# Estimating divergence times from DNA sequences

**DOI:** 10.1101/2020.10.16.342600

**Authors:** Per Sjödin, James McKenna, Mattias Jakobsson

## Abstract

The patterns of genetic variation within and among individuals and populations can be used to make inferences about the evolutionary forces that generated those patterns. Numerous population genetic approaches have been developed in order to infer evolutionary history. Here, we present the ‘Two-Two (TT)’ and the ‘Two-Two-outgroup (TTo)’ methods; two closely related approaches for estimating divergence time based in coalescent theory. They rely on sequence data from two haploid genomes (or a single diploid individual) from each of two populations. Under a simple population-divergence model, we derive the probabilities of the possible sample configurations. These probabilities form a set of equations that can be solved to obtain estimates of the model parameters, including population split-times, directly from the sequence data. This transparent and computationally efficient approach to infer population divergence time makes it possible to estimate time scaled in generations (assuming a mutation rate), and not as a compound parameter of genetic drift. Using simulations under a range of demographic scenarios, we show that the method is relatively robust to migration and that the TTo-method can alleviate biases that can appear from drastic ancestral population size changes. We illustrate the utility of the approaches with some examples, including estimating split times for pairs of human populations as well as providing further evidence for the complex relationship among Neandertals and Denisovans and their ancestors.

## Background

Many population genetic inference approaches compare levels of genetic variation within and across genomes, individuals and/or populations in order to uncover their evolutionary history. A multitude of demographic inference methods have been developed in order to capitalize on the wealth of information that comes with the availability of full genomes from multiple individuals (see Schraiber and Akey 2015, for a review).

The sheer scale and complexity of whole genome data sets poses its own challenge for making inference of population demographic parameters. A common approach for inference has been to compare the observed data, often summarized in some statistic, to simulated data that can be generated under a range of population-genetic models. Building on this idea and combined with a rejection algorithm, Approximate Bayesian Computation (ABC, Beaumont *et al.* 2002; Cornuet *et al.* 2014; Tavaré *et al.* 1997; Pudlo *et al.* 2016) has proven to be one useful tool for both model choice and parameter estimation. However, the problem of choosing which models to test is not trivial for most inference approaches, including ABC, as the set of models to choose from is very large.

In parallel, there have been recent developments in methods that use haplotype information. A challenge for these approaches is how to model the dependence of genealogies along a sequence. One solution has been to approximate the full ancestral recombination graph, a method used in PSMC (Li and Durbin 2011) and similar approaches (Schiffels and Durbin 2014; Terhorst *et al.* 2017; Kelleher *et al.* 2019; Speidel *et al.* 2019; Wang *et al.* 2020).

Another strategy has been to rely on relatively short genetic fragments located sufficiently far away from each other to be able to assume linkage equilibrium between loci, combined with absolute linkage (absence of recombination) within each locus (e.g. Gronau *et al.* 2011). Both these approaches typically lead to set-ups that cannot be solved analytically and often rely on computationally heavy, advanced statistical methods in order to estimate parameters (but see Gattepaille *et al.* 2016; Lohse *et al.* 2016). A related strategy is to assume independence among sites, using a composite likelihood framework (Gutenkunst *et al.* 2009; Excoffier *et al.* 2013). From this assumption, the observed variables (*i.e.* frequency spectra) do not depend on the full distribution of genealogical branch lengths, they are functions only of the expected branch lengths (Griffiths and Tavaré 1998; Chen 2012). This observation greatly simplifies the probability computations. To the extent that closed-form solutions can be obtained, the assumption of independence between sites also leads to inference tools that are easier to integrate with other methods, and can provide useful insights into underlying processes (Beichman *et al.* 2017; Terhorst *et al.* 2017). Conversely, a disadvantage of assuming independence between sites is that only information concerning the expected values can be obtained, rather than the full distributions of stochastic variables. For small samples, an alternative is to derive closed form expressions for the probability of observing particular configurations of variants in simple divergence models, including the isolation-with-migration model (Wakeley and Hey 1997; Wilkinson-Herbots 2008; Chen 2012). Lohse *et al.* (2011, 2016) showed that more generally, the probability of observing a particular variant configuration can be obtained from a generating function of genealogical branches. Assuming independence among sequence-blocks, Lohse *et al.* (2011, 2016) outlined an approach for computing the likelihood under various demographic models and sampling schemes.

Regardless of whether independence between sites is assumed or not, all of these methods can be useful for inferring the time of divergence between two populations. Examples of simple and direct methods used to estimate population divergence times include: Green *et al.* (2010); Gutenkunst *et al.* (2009); Wakeley (2009); Schlebusch *et al.* (2012); Theunert and Slatkin (2018). These methods build on the principle of genetic drift accumulating as a function of effective population size and number of generations. Following a population backwards in time, and using that the accumulated drift at generation *t* is,

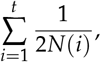

where *N*(*i*) is the (effective) number of individuals at generation *i*, the divergence time is then the number of generations required to generate the estimated drift. Such estimates are typically not dependent on knowing the mutation rate but some assumptions regarding *N*(*i*) is required, either by assuming a fixed effective population size or depending on an estimated function of *N*(*i*).

Alternatively, one can base the divergence time estimate on an assumed mutation rate (e.g. Wakeley and Hey 1997; Chen 2012; Pickrell *et al.* 2012). By assuming independence among sites, in a two-population divergence model (without migration), the probability of observed sample configurations (summarized as the full SFS, including invariable sites) can be derived analytically. Using a likelihood framework, we can then estimate parameters of interest in the divergence model. Here we present two simple approaches based on picking two gene copies from each of two populations: the ‘Two-Two’ (TT) method, which was briefly introduced in Schlebusch *et al.* (2017) and the ‘Two-Two-outgroup’ (TTo) method. These are sufficiently simple to allow for analytical solutions giving closed formulas for the estimates of the model parameters based on the counts of different sample configurations. Specifically, we can directly estimate population divergence time in generations, an estimate which is independent of genetic drift and effects of varying population size since the model is parametrized with both a drift parameter and a time parameter.

## Observed data

For the purpose of investigating the demographic relationship between two populations denoted population 1 and population 2, assume that two gene copies have been sampled from each population. For bi-allelic sites, assume that the ancestral (denoted ‘0’) and the derived variant (denoted ‘1’) is known. The number of derived alleles in a sample from population 1 combined with the number of derived alleles in as sample from population 2 is referred to as the joint frequency spectra. Following Chen (2012), a sample configuration of *k*_1_ derived variants in a sample of size *n*_1_ from population 1 and *k*_2_ derived variants in a sample of size *n*_2_ from population 2 can then be denoted as *S*_*n*1,*n*2_(*k*_1_, *k*_2_). In our set up, *n*_1_ = *n*_2_ = 2 and *k*_1_, *k*_2_ are either 0, 1 or 2, and there are 9 possible sample configurations, which are presented in Table 1. The observed number of sites with sample configuration *O*_*i,j*_ will be denoted by *m*_*i,j*_ and the total number of investigated sites by *m*_*tot*_.

**Table 1.**
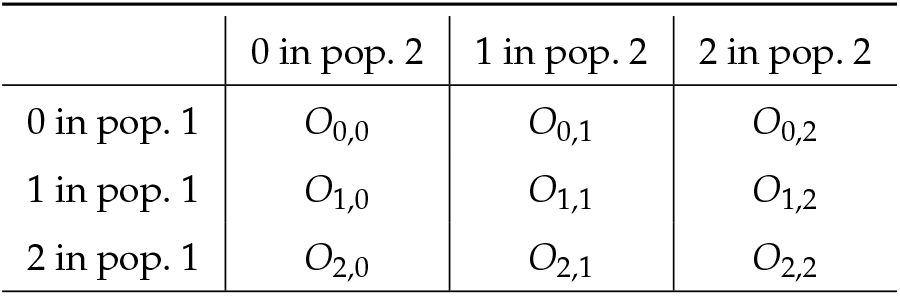
Notation for the number of sites with 0, 1, or 2 derived variants in the sample from population 1 and the sample from population 2.

## Theory

We study a general population divergence model where the population-branch leading to population 1 and the population-branch leading to population 2 merge (backwards in time) to become the ancestral population. The model makes no assumptions regarding population size and/or population structure changes in the daughter populations. The model assumes no migration between the two daughter populations and that these merge into a panmictic ancestral population.

We use the following notation, with time measured in number of generations:

*t*_1_ : time to split for pop. 1,
*t*_2_ : time to split for pop. 2,
*α*_1_ : prob. of two lineages in pop. 1 not coalescing before *t*_1_,
*α*_2_ : prob. of two lineages in pop. 2 not coalescing before *t*_2_,
*ν*_1_ : expected time to coal. in pop. 1 given coal. before *t*_1_,
*ν*_2_ : expected time to coal. in pop. 2 given coal. before *t*_2_.

In addition to the drift parameters *α*_1_ and *α*_2_, the parameters *ν*_1_ and *ν*_2_ are needed because two branches with the same time-length and the same drift can have different distributions of coalescence times. To illustrate, a linearly growing population that starts with size *N* and ends with size 2*N* will have the same drift as a shrinking population that starts with size 2*N* and ends with size *N* but they will not have the same distribution of coalescent times within that interval. These parameters also cover cases when the daughter populations are not panmictic. A similar parametrization can be found, for instance, in Rogers and Bohlender (2014).

The composite likelihood assumption of independence between sites implies that the probability of a mutation on a specific branch in a genealogy is the expected length of that branch (given a demographic model) multiplied by the mutation rate. We denote the mutation rate per site and generation by *μ*, assume independence between sites, and an infinite sites model.

We define the following events for the two sampled lineages for each population:

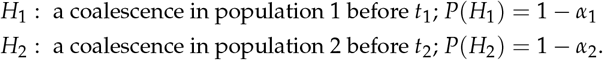

With 2 ≤ *k ≤* 4 lineages surviving to enter the ancestral population (depending on whether coalescence events have occurred in the daughter populations), we define *A*_*k*_ to be the number of derived variants in a sample of size *k* drawn at the split time in the ancestral population and write *a*_*ki*_ = *P*(*A*_*k*_ = *i*). To illustrate how the probabilities of the sample configurations are derived, we can take an example conditional on no coalescent event in population 1 and a coalescent event in population 2 (the event ¬*H*_1_ ∧ *H*_2_). There are then 3 lineages entering the ancestral population. These lineages constitute a sample of size 3 from the ancestral population. Sample configuration *O*_1,0_ will then be observed with probability (2/3)*a*_31_ (the ancestral variant has to be assigned to the lineage entering population 2; an event with probability 2/3) plus the probability that a mutation occurs on either lineage entering population 1 during the time interval *t*_1_. The probability that a mutation hits a branch of length *t*_1_ is *μt*_1_, and the probability that this happens *and* that the derived variant already exists in the ancestral population can be ignored as it requires two mutational events at the same site. Thus, conditional on ¬*H*_1_ ∧ *H*_2_, *P*(*O*_1,0_) = (2/3)*a*_31_ + 2*μt*_1_. The same reasoning can be applied to derive the conditional probabilities for all 7 (polymorphic) sample configurations and these are shown in Table 2.

**Table 2.** Conditional probabilities.

| | $H_1 \wedge H_2$ | $H_1 \wedge \neg H_2$ | $\neg H_1 \wedge H_2$ | $\neg H_1 \wedge \neg H_2$ |
| --- | --- | --- | --- | --- |
| $O_{1,0}$ | $2\mu\nu_1$ | $2\mu\nu_1$ | $\frac{2a_{31}}{3} + 2\mu t_1$ | $\frac{a_{41}}{2} + 2\mu t_1$ |
| $O_{0,1}$ | $2\mu\nu_2$ | $\frac{2a_{31}}{3} + 2\mu t_2$ | $2\mu\nu_2$ | $\frac{a_{41}}{2} + 2\mu t_2$ |
| $O_{2,0}$ | $\frac{a_{21}}{2} + \mu(t_1 - \nu_1)$ | $\frac{a_{31}}{3} + \mu(t_1 - \nu_1)$ | $\frac{a_{32}}{3}$ | $\frac{a_{42}}{6}$ |
| $O_{0,2}$ | $\frac{a_{21}}{2} + \mu(t_2 - \nu_2)$ | $\frac{a_{32}}{3}$ | $\frac{a_{31}}{3} + \mu(t_2 - \nu_2)$ | $\frac{a_{42}}{6}$ |
| $O_{1,1}$ | 0 | 0 | 0 | $\frac{2a_{42}}{3}$ |
| $O_{2,1}$ | 0 | $\frac{2a_{32}}{3}$ | 0 | $\frac{a_{43}}{2}$ |
| $O_{1,2}$ | 0 | 0 | $\frac{2a_{32}}{3}$ | $\frac{a_{43}}{2}$ |

Since a subsample of size *n* randomly drawn from a larger sample of size *n* + *k* has the same distribution as a sample of size *n* drawn directly from the population, we can reduce the number of parameters by replacing all *a*_*ij*_ with *i <* 4 using *a*_*ij*_-terms with *i* = 4 as follows:

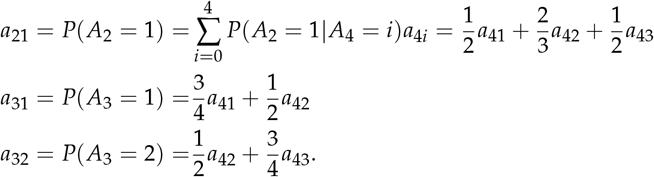

These equations together with Table 2 allows us to derive the probabilities for the different sample configurations. For instance:

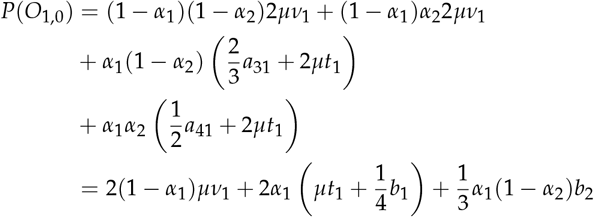

where *b*_*i*_ = *a*_4*i*_ = *P*(*A*_4_ = *i*).

Using the same strategy for the derivation of the other six probabilities, we obtain the probabilities for all seven sample configurations in Table 2. Writing *p*_*i,j*_ = *P*(*O*_*i,j*_) for brevity, these are

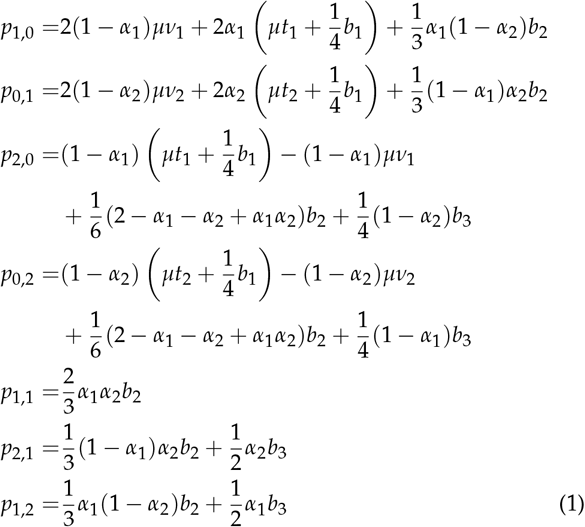

Furthermore, if we assume a (indefinitely) panmictic ancestral population (figure 1a), we define

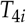

to be the number of generations a coalescent process that starts with 4 lineages at the (most recent) base of the ancestral population spends with *i* lineages, so that *T*_*mrca*_ = *T*_44_ + *T*_43_ + *T*_42_. Then (see appendix)

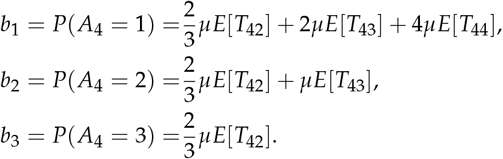

**Figure 1.**
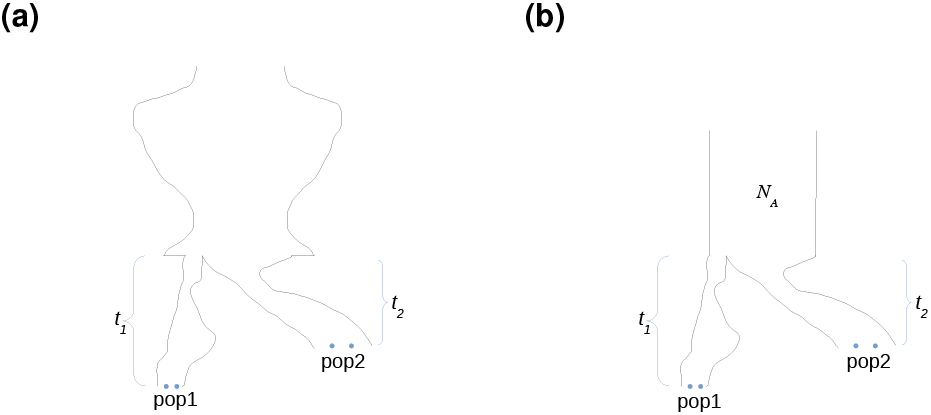
Different assumptions for population divergence models (a) panmictic ancestral population, and (b) constant ancestral population.

Writing *τ*_*i*_ = *μE*[*T*_4*i*_], and replacing the *b*_*i*_ with their respective expression in terms of *τ*_*i*_, the probabilities for the different sample configurations can be expressed as:

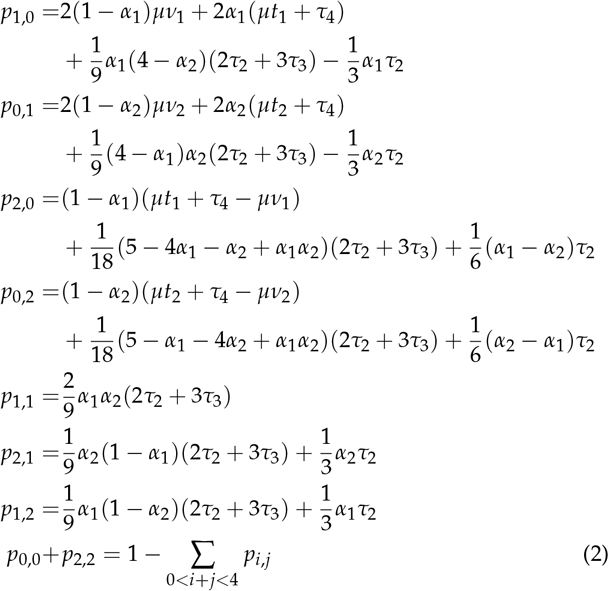

These 8 equations point to two challenges: i) it is not possible to completely separate *τ*_4_ from divergence times due to its co-occurrence with *μt*_1_ and *μt*_2_, ii) disregarding *τ*_4_, it is still an under-determined set of equations with 8 parameters but only 7 equations/degrees of freedom (*p*_0,0_ + *p*_2,2_ = 1 − ∑_0*<i*+*j<*4_ *p*_*i,j*_). It can be tempting to reduce the number of parameters by setting *t*_1_ = *t*_2_, but because (from equations 1 above)

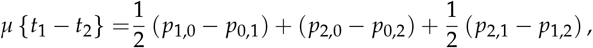

specifying *t*_1_ = *t*_2_ would add additional dependence between the equations. Although this will decrease the number of parameters, it also decreases the number of independent equations. Furthermore, allowing for separate divergence times along the two branches is a valuable asset; not only does it allow the framework to be applicable for temporally structured samples, but separate estimates for each branch can be useful more generally. In fact, it turns out that the divergence time estimate based on the population branch represented by a modern-day individual alleviates the potential issue of residual ancient DNA specific properties (DNA degradation, sequencing errors, mapping errors) that could impact divergence time estimates (see below). In contrast, for contemporaneous samples, divergence time estimates should be the same along the two branches (assuming neutrality and the same mutation rate and generation time along the two branches).

The challenges noted above can be dealt with either by assuming a constant ancestral population size (the “TT”-method) or by using an outgroup to increase the number of equations (the “TTo”-method).

### Assuming a constant ancestral population size (’TT’)

Assuming a constant ancestral population size *N*_*A*_ reduces the number of parameters in the model (figure 1b), so that *E*[*T*_4*k*_] = 2*N*_*A*_/(*k*(*k* − 1)) and (with *θ* = *μN*_*A*_) *τ*_2_ = *θ*, *τ*_3_ = *θ*/3 and *θ* = *τ*_4_/6. Then the probabilities in equations (2) simplify as

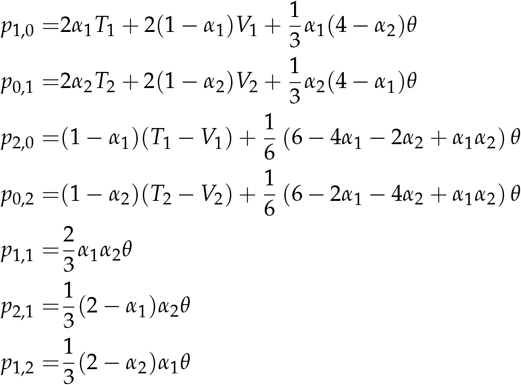

With

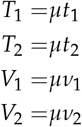

We set 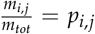 and solve for the parameters (and note that this guarantees that they are also maximum likelihood estimates (Doob 1934; Wald 1949) to obtain:

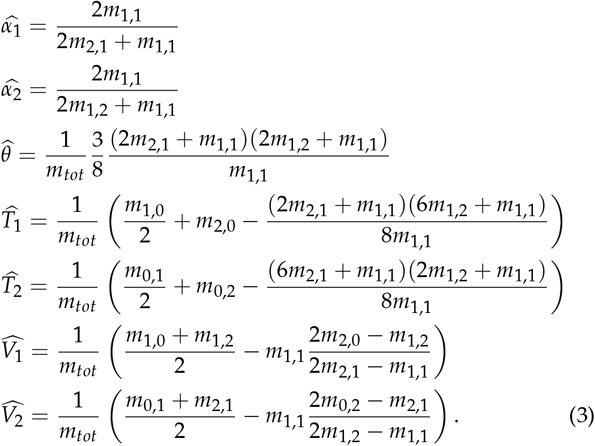

These equations are referred to as the ‘TT’-method in Schlebusch *et al.* (2017) where, in order to get the divergence time in years we used *G* = 30 and *μ* = 1.25 × 10^−8^ in

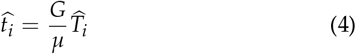

where *G* is the length of a generation.

Note also that if sequencing errors or DNA-degradation mainly result in additional singletons, then errors in the sample from population 1 only affects *m*_1,0_ and thus only 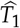 and 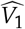 (*m*_1,0_ occurs exclusively in the equations for estimating *T*_1_ and *V*_1_).

### Adding an outgroup (’TTo’)

The equations (1) are useful for data (including SNP-genotype data) where the derived variant at each site has been ascertained in a population that branched off prior to the investigated population split. Such data will ensure that derived variants in the studied sample will be older than the split so that there are no new mutations occurring in the branches. In such a case, *μt*_1_, *μt*_2_, *μν*_1_ and *μν*_2_ can all be set to 0 in the equations 1 above, resulting in a new set of equations (see appendix) that can be solved for the *α*’s to get

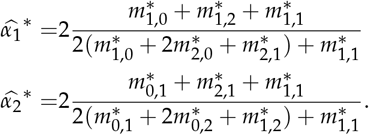

where * indicates that these are the corresponding parameters and sample configuration counts conditional on ascertainment in an outgroup.

With this ascertainment procedure it is important that the population used to ascertain the SNPs represents a true outgroup to our studied populations and that the populations satisfy an assumption of bifurcating topology (or “tree-ness”). To validate such an assumption we can set up tests of tree-ness, since if *μt*_1_ = *μt*_2_ = *μν*_1_ = *μν*_2_ = 0, then the test statistics

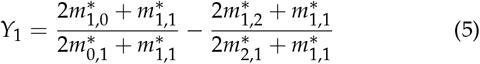

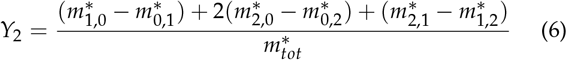

should be 0 (see appendix, where it is also shown that *Y*_2_ is closely related to the D-statistic, Green *et al.* 2010).

The estimates 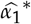 and 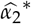 of *α*_1_ and *α*_2_ together with the equations in (2) can furthermore be used to obtain estimates of

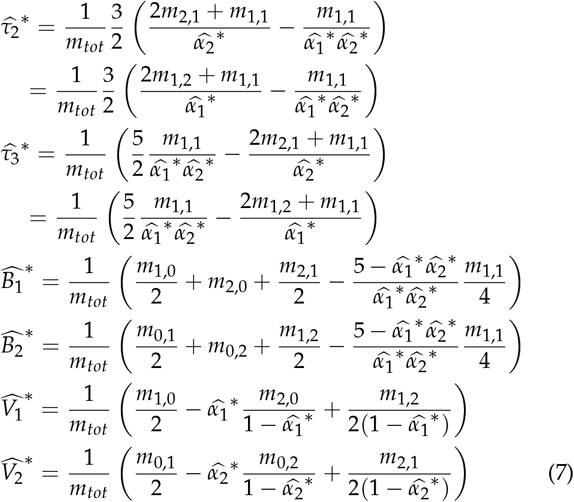

with

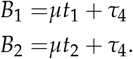

Note the two alternative estimates (one using *m*_2,1_ and one using *m*_1,2_) for *τ*_3_ and *τ*_2_ and we take the average of these in estimates below.

Based on the obtained estimates of *τ*_2_ and *τ*_3_, we can attempt to approximate *τ*_4_ as a combination of *τ*_2_ and 3*τ*_3_. In a constant population, *E*[*T*_43_]/*E*[*T*_42_] = 1/3 and *E*[*T*_44_]/*E*[*T*_43_] = 1/2 or

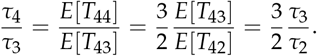

For this reason we propose to approximate *τ*_4_/*τ*_3_ as (3/2)*x* where *x* is the estimated ratio of *τ*_3_/*τ*_2_. This leads to

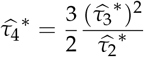

to get

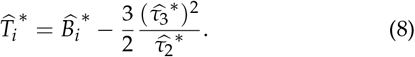

We refer to this approach to estimate divergence time as “TTo” (as in “TT outgroup”).

### Picking two gene copies from population 1 and one gene copy from population 2

The method so far described can be seen as an expansion of the simpler case of picking two gene copies from one population, and only one gene copy from the other population. This simpler set-up can be useful, for instance, when dealing with low coverage genome data (e.g. ancient DNA sequence data). With this simpler approach, divergence time estimation needs an out-group (only assuming a constant population size is not sufficient to solve the equations in this case). This 2 plus 1 approach does, however, provide reliable estimates of branch specific genetic drift (under often reasonable demographic assumptions, see appendix and Wakeley 2009; Schlebusch *et al.* 2012; Skoglund *et al.* 2011).

### Simulations and comparison to GPhoCS

The model underlying the TT method assumes a panmictic ancestral population of constant size prior to the split, and no gene-flow between populations after the split. While common to many coalescent-based approaches, such assumptions are rarely realistic for natural populations, and it is increasingly evident that mis-specification of an overly simplistic model may lead to substantially biased parameter estimates (Gronau *et al.* 2011; Mazet *et al.* 2016; Orozco 2016).

Here we investigate the robustness of the TT-method parameter estimation against violation of the basic model assumptions (equations in 3). We compare its performance under these conditions against an alternative method for parameter inference, GPhoCS (Gronau *et al.* 2011). The analytical TT method and the Bayesian inference method GPhoCS are located to some degree at opposite ends of the statistical inference spectrum; instead of relying on independent single bi-allelic sites, GPhoCS assumes complete linkage between individual sites at a genetic locus (typically 10 kb), but independence between these loci. It should be noted that GPhoCS is capable of estimating parameters under more complicated demographic models than the simple split model we study here. In particular, GPhoCS allows users to specify migration rates and define migration bands between populations, such that it does not share the TT method assumption of no gene-flow occurring between populations after the population split. For this reason, the effect of migration on parameter estimation was investigated only for the TT method.

The software MS (Hudson 2002) was used to generate polymorphic datasets using a standard coalescent algorithm under a variety of demographic scenarios. The effects of changes in ancestral population size (figure 2A) and migration between branches since the split (figure 2B) were investigated. In each model, the ancestral population size, *N*_*A*_, was fixed at 34,000, corresponding to 17,000 diploid individuals. This value is in line with recent estimates of African ancestral effective population size approximately 1 million years ago (Li and Durbin 2011; Schiffels and Durbin 2014; Schlebusch *et al.* 2017). MS scales time by 4*N*_*e*_, and simulations were constructed with true split times of 10,000 and 1500 generations. Assuming a generation time of 30 years this equates to split times of 300,000 and 45,000 years respectively. These were chosen to keep simulations relevant to the findings of previous work where the deepest split among human groups was estimated at >260,000 years (Schlebusch *et al.* 2017), together with more recent divergence events.

**Figure 2.**
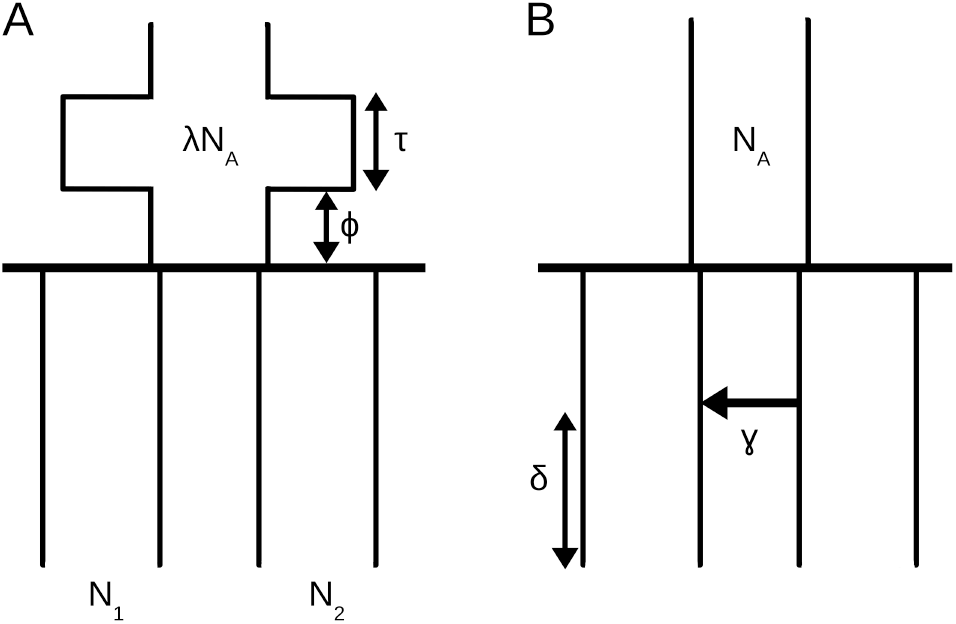
The two general demographic models used to simulate data for testing robustness of TT method, with (A) changes in ancestral population size, and (B) variation in proportion and timing of migration between branches since a population split.

MS generates samples assuming *θ* = 4*N*_1_*μ*, where *N*_1_ is the diploid population size of population 1, and *μ* is the neutral mutation rate. Mutation rates vary across the human genome and estimates vary depending on the method used (Scally and Durbin 2012). Li and Durbin (2011) calculated a human neutral mutation rate of 2.5 × 10^−8^ per generation (assuming 25 years per generation), whilst recent consensus suggests a lower rate of 1.25 × 10^−8^ per base pair per generation is more accurate (Moorjani *et al.* 2016). The latter is the mutation rate used across all simulations. Results were filtered such that only those simulations resulting in all sample configurations represented by > 10, 000 sites were used in subsequent analyses.

The Bayesian inference method GPhoCS is based on likelihood estimation and in order to allow adequate convergence of parameter estimates, a burn-in period of 100,000 iterations was used when applied to MS simulated data.

### The effect of varying ancestral population size

In simulating the demographic scenario shown in figure 2A, populations 1 and 2 are constant backwards in time, but not (necessarily) equal in size; each population size is independently drawn from a uniform distribution between 170 and 1,700,000 diploid individuals. A total of 1000 of such demographies were generated. Populations 1 and 2 merge at 10,000 generations to form a single ancestral population of initial size *N*_*A*_ = 17, 000 individuals for *ϕ* generations. The ancestral population then changes to *λN*_*A*_, (drawn uniformly from [1.7 × 10^2^, 1.7 × 10^6^]), for *τ* generations, before returning to *N*_*A*_. We investigate the impact of that change in ancestral population size (*λN*_*A*_ for *τ* generations) on TT estimates of population divergence time 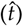 and ancestral population size, 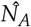, for *τ* = 0, 100, 500, 1, 000 generations and *ϕ* = 0, 100, 500, 2, 000 generations (figure S1). Figure S1 shows that increasing ancestral population size for *τ* generations can have the effect of inflating divergence time estimates for the TT method. This behaviour is expected; if we imagine an expansion of the ancestral population to infinite size for *τ* generations, no coalescence events would occur during that time, and divergence time estimates would be upwardly biased by *τ* generations. This bias however seems to be relatively minor compared to that arising from severe bottlenecks. For instance, when the true *t* = 10, 000, a bottleneck in the ancestral population of 1,500 individuals lasting for 100 generations results in 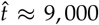 generations. However, the same severity of bottleneck lasting for 500 generations will result in a greater underestimate of 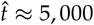 generations. Similarly, *N*_*A*_ is under-estimated when severe bottlenecks occur, though these estimates seem to be more robust than estimates of divergence time (figure S2). When studying more recent splits, (1500 generations), we observe that severe bottlenecks have the potential to result in nonsensical negative split time estimates (figure S3).

Figure 3 shows a comparison between TT method and GPhoCS estimates of population divergence time (*t*) and ancestral population size (*N*_*A*_) in cases where the duration of the bottleneck (*τ*) is fixed at 1,000 generations and true split time is 10,000 generations. Results suggest that both methods react similarly to violations of the assumption of a change in ancestral population size; each being particularly susceptible to bias when severe bottlenecks have occurred. GPhoCS performs somewhat better than the TT method, with severe bottlenecks resulting in less of an underestimate of population divergence time.

**Figure 3.**
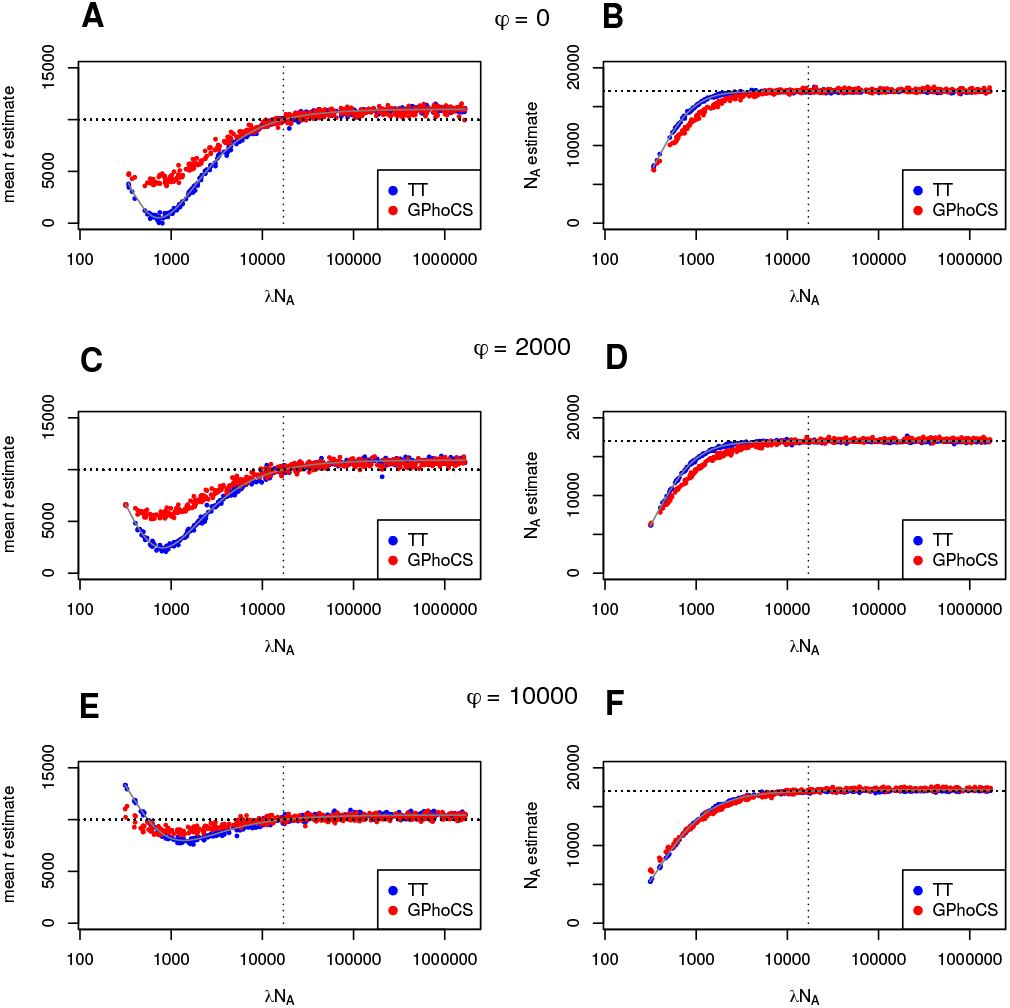
A comparison of the effect of ancestral population size changes on TT method and GPhoCS parameter estimates. The time between population divergence and change in ancestral population size (*ϕ*) is (A, B) 0, (C, D) 2,000, and (E, F) 10,000 generations. In all cases, the duration of change in ancestral population size (*τ*) is 1,000 generations and true split time is 10,000 generations.

An interesting effect appears when a bottleneck of sufficient severity occurs, whereby both methods’ 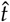 estimates begin to rebound towards the true split time of 10,000 generations. Again this behaviour is expected as all lineages will coalesce in a bottleneck of sufficient severity prior to a population divergence event. In this case the bottleneck itself will act as the constant ancestral population size, and as long as it occurs in close proximity to the split, divergence time estimates are not affected much. For the same reason, when a severe bottleneck occurs a long time prior to the split, both methods produce a (slight) overestimate of the true divergence time (figure 3E).

### The effect of migration between branches

In simulations based on the demographic scenario shown in figure 2B, a pulse of admixture occurs *δ* generations ago, with proportion 0 ≤ *γ* ≤ 1 of one daughter population made up of migrants from the other daughter population. Thus we examine the effect of increasing proportion of migration occurring at various times between present and the split time. All populations are kept fixed and constant at 17,000 diploid individuals (*N*_1_ = *N*_2_ = *N*_*A*_). Figures 4 and 5 show the effect of increasing proportion of migrants on TT divergence time estimates when true split time is 10,000 generations. Divergence times are reliably estimated when the proportion of migrants is below 0.1, and as expected, even at higher proportions the bias decreases the nearer the admixture event is to split time. Note that under this set up, the TT method returns 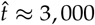 generations even when the proportion of migrants (*γ*) is 1 and admixture time (*δ*) is 0. We would expect in this case a 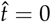, but it seems that this scenario (where all populations are equal in size) is equivalent to a violation of the assumption of a constant ancestral population. As described previously, this has the effect of biasing 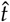 upwards, and shows that in cases of high proportion of recent admixture, differences between the size of the daughter populations and the ancestral population also has the potential to result in biased 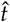. Estimates of ancestral population size on the other hand, are only very slightly affected (figures S4 and S5). Very similar results are observed when a more recent true split time of 1500 generations is studied (figures S6 and S7).

**Figure 4.**
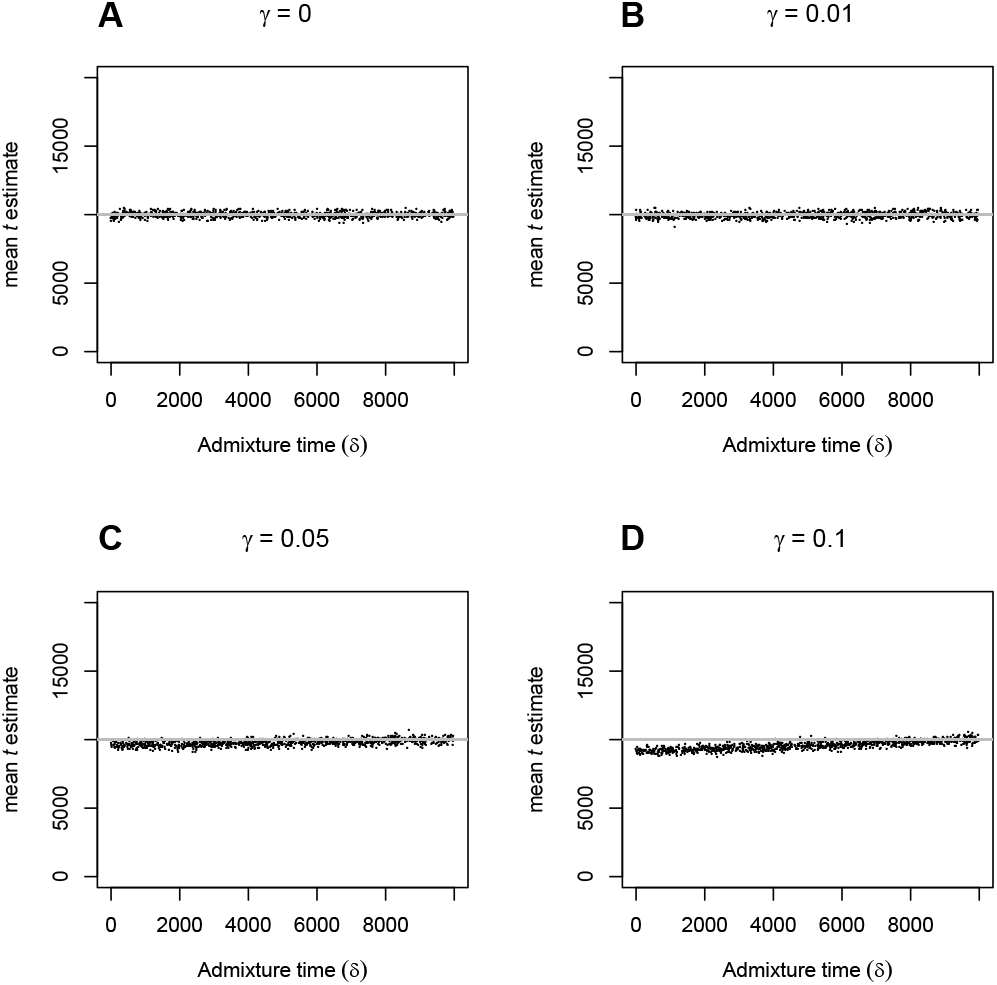
The effect of varying admixture time (*δ*) on TT split time estimates 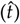, when proportion of admixture (*γ*) is (A) 0, (B) 0.01, (C) 0.05 and (D) 0.1, and true split time is 10,000 generations.

**Figure 5.**
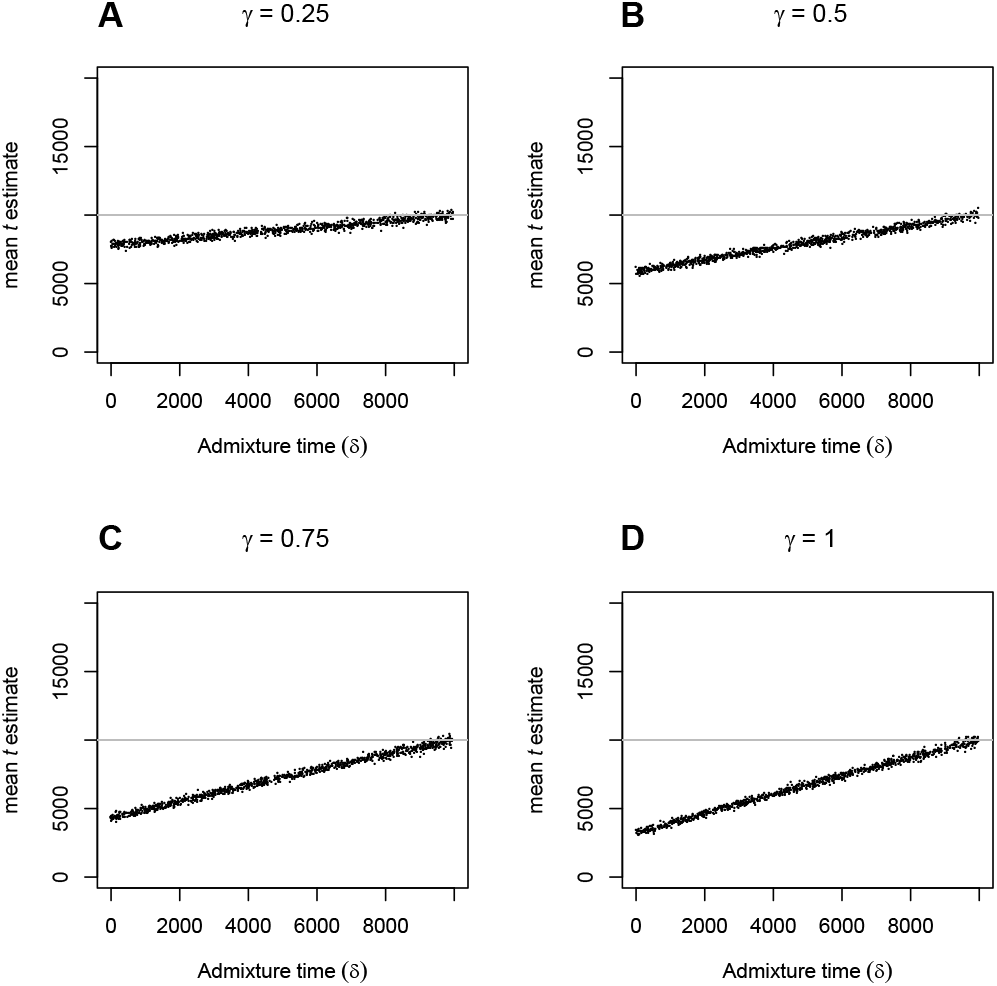
The effect of varying admixture time (*δ*) on TT split time estimates 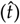, when proportion of admixture (*γ*) is (A) 0.25, (B) 0.5, (C) 0.75 and (D) 1, and true split time is 10,000 generations.

While simulations have shown the TT method to be relatively robust to violations of its assumptions in general, it is evident that extensive, recent gene flow between daughter populations or strong, prolonged bottlenecks in the ancestral population have the potential to introduce bias. If however we obtain external estimates of *α*_1_ and *α*_2_ through the outgroup ascertainment procedure outlined above (TTo), we can obtain estimates of divergence time that are much less dependent on assumptions concerning the ancestral population. Figure 6 shows a comparison of TT and TTo method results in scenarios of increasing duration of ancestral population size change. The true values of *α*_1_ and *α*_2_ have been used in equations in 7 to obtain estimates of *B*_1_, *B*_2_, *τ*_2_ and *τ*_3_, that in turn have been used to approximate *τ*_4_ and divergence times following equation 8. These results show that by using external estimates of drift, there is the potential to considerably reduce bias in divergence time estimates when severe bottlenecks have occurred in the ancestral population. Furthermore, figure 6 also compares estimates of *N*_*A*_, from that of the TT method to one based on the TTo estimate of *τ*_4_ 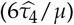, which is found to be much more sensitive to ancestral bottlenecks.

**Figure 6.**
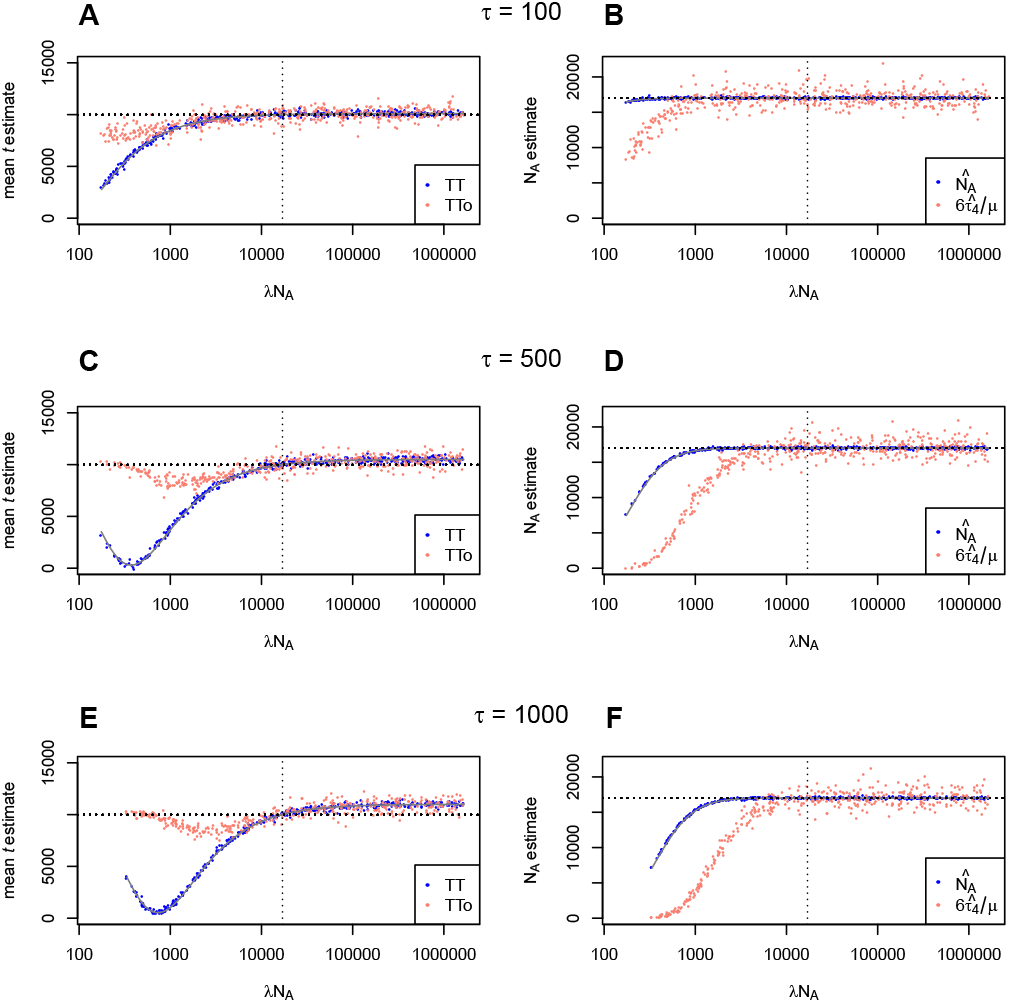
A comparison of TT method estimates of divergence time and ancestral population size with (TTo), and without (TT) using external estimates of drift. The duration of alternative ancestral population size (*τ*) is (A, B) 100, (C, D) 500, and (E, F) 1,000 generations. In all cases, the change in ancestral population size occurs immediately prior to the split (*ϕ*=0) and true split time is 10,000 generations.

## Application to data

The TT method requires good quality sequence data, typically high-coverage genome sequence data since diploid genotype calls are utilized, including invariable sites and singletons (in the sample of four chromosomes). The formulas are sufficiently simple to allow for asymptotic confidence intervals based on MLE theory, and one can imagine thinning the genome data to make sites independent of each other (to overcome potential dependence via linkage). However, we chose a more conservative approach of estimating confidence; the weighted block jackknife procedure (Busing *et al.* 1999), which should be more robust to large-scale “outlier” regions driving the signal. We conducted pairwise comparisons among the 11 HGDP individuals and the Denisovan genome and the Altai Neandertal genome from (Meyer *et al.* 2012) and (Prüfer *et al.* 2014) and estimate population divergence times (see appendix for a description of data cleaning and calling of ancestral states). The 11 HGDP individuals include 5 individuals from Africa: one Khoe-San (‘San’), one rainforest hunter-gatherer (‘Mbuti’), two West-African (‘Mandenka’ and ‘Yoruba’) and one East-African (‘Dinka’). The other individuals were two Europeans (‘French’ and ‘Sardinian’), two East-Asians (‘Han’ and ‘Dai’), one individual from Oceania (‘Papuan’) and one individual representing a South-American indigenous population (‘Karitiana’). In addition, we used the high coverage ancient southern African hunter-gatherer genome (‘Balito Bay A’, (Schlebusch *et al.* 2017) as an outgroup for some divergence estimates (see below).

### Split model parameter estimates

We refer to split time estimates *in years* in the TT-method as 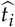 and under the TTo-method as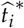. These are obtained by setting *G* = 30 and *μ* = 1.25 × 10^−8^ in equation (4) and applying this to the estimates of *T*_1_ and *T*_2_ in equations (3) and equation (8) respectively.

Comparisons are grouped according to the population split they represent. For instance, the comparison between French and San is referred to as the ‘Khoe-San split’.

### Divergence estimates according to the TT-method

Assuming a constant ancestral population size, *N*_*A*_, it is possible to estimate *N*_*A*_, *α*_1_, *α*_2_, *ν*_1_, *ν*_2_ as well as *t*_1_, *t*_2_ without relying on ascertainment procedures. Estimates of *α*, *N*_*A*_ and *ν* are shown in supplementary figures S9, S8 and S10 respectively. From figure S10 it is apparent that *ν* is often poorly estimated and the uncertainty of the estimate appears to be closely linked to the amount of branch-specific genetic drift (figure S11). A closer look at any of the formulas for *p*_*i,j*_ reveals that the impact of *ν* on the probabilities disappears as *α* approaches 1 (no drift).

Estimates of the ancestral population size remain remarkably constant at around *N*_*A*_ = 17, 000, regardless of choice of individuals (figure S8).

Values of 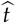 are shown in figure 7. To summarize, estimates of the different split times are (in descending order)

- the split between Neanderthal and Denisovans 962 − 979 kya;
- the split between archaic humans and modern humans 510 – 707 kya;
- the deepest split among modern human population (between Khoe-San and other human populations) 233 − 266 kya (see (Schlebusch *et al.* 2017) for the consequence of using the ancient southern African Balito Bay A genome);
- the split between Mbuti and other modern humans (excluding Khoe-San populations) 186 – 220 kya
- the split between West-Africans and East-Africans 96 – 117 kya;
- the split between East-Africans and non-African 66 – 82 kya;
- splits between non-African < 0 ya.

**Figure 7.**
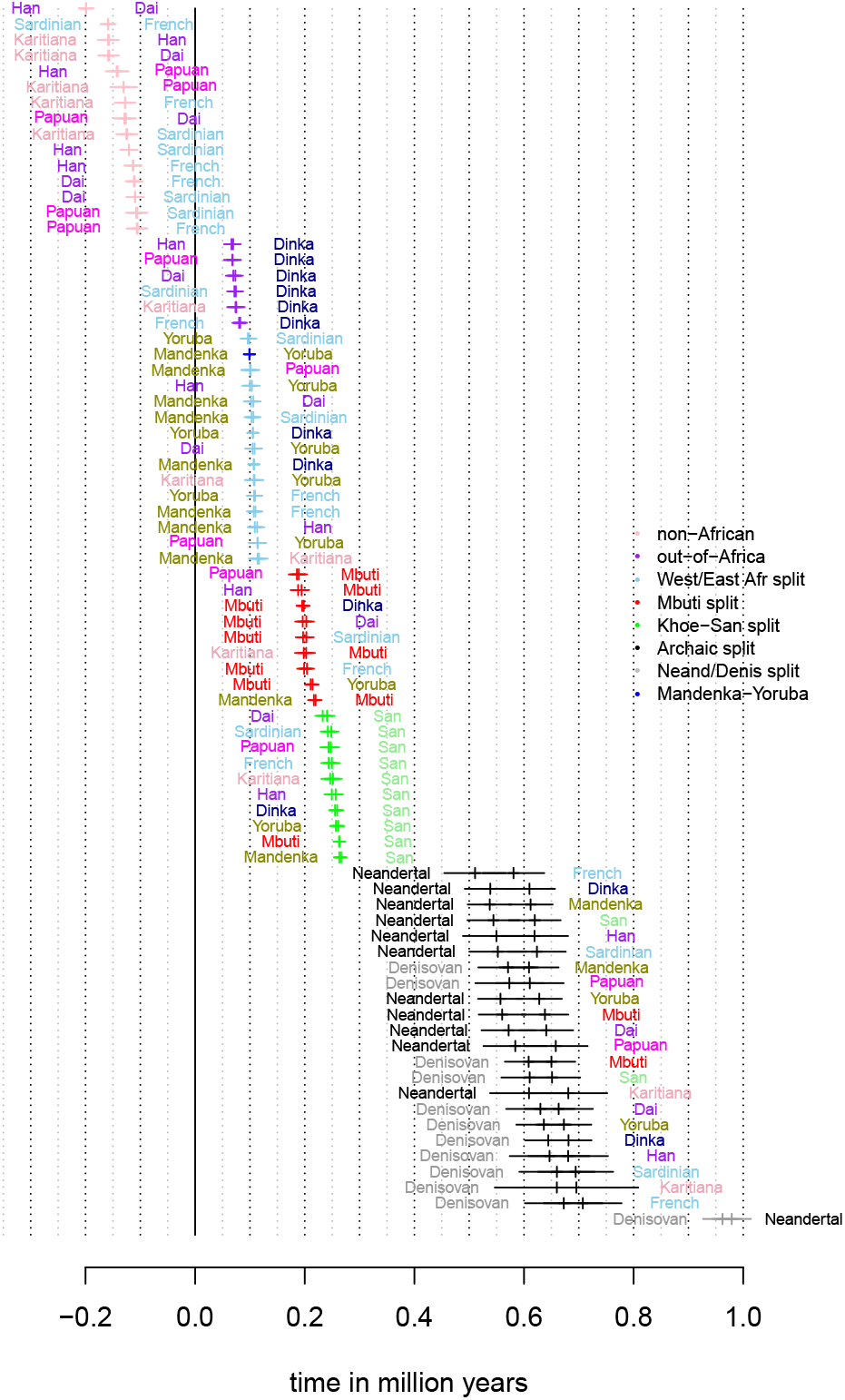
Split time estimates assuming a constant ancestral population and a mutation rate of 1.25 × 10^−8^ and a generation time of 30 years. Corresponding to the branch specific divergence time, there are two estimates each comparison.

Here, the range for the split between archaic and modern humans takes into account the fact that the archaic genomes are older than 40 ky. There are two obvious odd sets of estimates among these: the negative times for non-Africans, and the deep time between Denisovans and Neandertals contrasted to the younger time between Denisovans/Neandertals and modern humans (note that we assume a constant ancestral population size here). We discuss each of these split time estimates below, but first we revisit the utility of ascertaining variants in an outgroup.

### Divergence estimates according to the TTo-method

By comparing two individuals using only those sites where the derived variant was present in an outgroup, it is possible to: i) test whether the outgroup represents a true outgroup, and ii) obtain estimates of *α*_1_ and *α*_2_ that do not rely on assumptions concerning the ancestral population. We utilized the Mbuti, Balito Bay A or Neandertal/Denisovan as outgroups. The estimates of *α* conditional on the derived variant being present in an outgroup are shown in figure S15. These three options were variably suitable as outgroups depending on the comparison being made. For instance, when comparing an individual from outside Africa to an African individual, Neandertal/Denisovan would not be true outgroups given the archaic admixture shared among non-African individuals (Green *et al.* 2010). This was also visible in the tests based on equations 5 and 6 above (as well as the D-test, see figure S12 in supplementary material). A likely consequence of the documented additional Denisovan ancestry in Papuan (Meyer *et al.* 2012) is that no comparison involving Papuan passed the outgroup tests. Perhaps more surprising, any comparison involving Mbuti failed the tests when Balito Bay A was used as the outgroup. Moreover, both Mbuti and Balito Bay A were expected to be true outgroups for the comparison of Neandertal versus Denisovan, but the test, however, pointed to them not being true outgroups.

Comparisons between estimates of *θ* (assuming a constant ancestral population size) to estimates of *τ*_2_ and 3*τ*_3_ using different outgroups for ascertainment are shown in figures S16, S18 and 17. Since there is presently no suitable outgroup for comparisons between a modern human and one of the two archaic humans – this would require a genome from an archaic human that split off before the Neandertal/Denisovan branch – it was not possible to estimate *τ*_2_ and 3*τ*_3_ for such comparisons.

When reliable outgroup ascertained estimates of *α*_1_ and *α*_2_ can be obtained, we estimate *τ*_2_, *τ*_3_, *B*_1_ and *B*_2_ using equations 7 that are used in equation (8) to obtain an estimate of 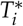. This in turn gives 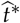 that are shown in figures S16, S17, S18, S19, S20, S21. For the majority of comparisons, such an approach does not yield different estimates compared to assuming a constant ancestral population size. The major exceptions to this are those comparisons involving non-Africans, that shows positive and realistic divergence time estimates using the ascertainment scheme (Figure 8).

**Figure 8.**
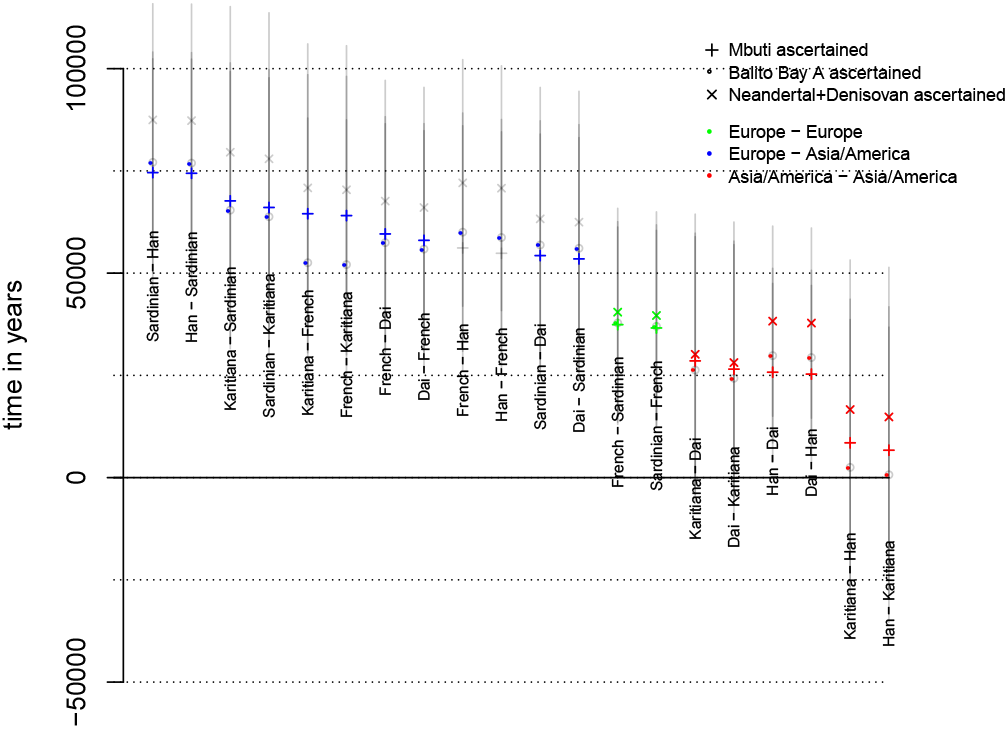
Different estimates of split times using outgroup ascertainment assuming a mutation rate of 1.25 × 10^−8^ and a generation time of 30 years. Comparisons with Papuans not included as no such comparison passed the outgroup-tests. Three estimates are shown: estimates where outgroup ascertainment is performed in Mbuti (+), in Balito Bay A (o) and in Neandertal/Denisovan (×). Transparent grey represents standard deviation and for comparisons that failed the outgroup tests.

### Divergence times outside Africa

The divergence time estimates for non-African populations under a constant model 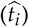 are nonsensical, (negative values). This is likely a consequence of the severe out-of-Africa bottleneck that leads to *τ*_4_ = *μE*[*T*_44_] being much smaller than *τ*_2_/6, which then violates the assumption of a constant *N*_*A*_(*E*[*T*_*nk*_] = 2*N*_*A*_/*k*(*k* − 1) in a constant population with *N*_*A*_ chromosomes). Estimates based on the three outgroup ascertainment schemes 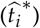 give more reasonable values as shown in figure 8.

Here, the split times estimates are:

- 50 to 75 kya between Europeans and Asians/Americans
- ~ 40 kya between Sardinians and French
- 25 to 30 kya between Dai and Karitiana
- 25 to 30 kya between Dai and Han
- < 25 kya between Han and Karitiana

These estimates are generally consistent with the prevailing view of the demographic history outside Africa. For instance, the deep split between Sardinians and French may reflect previous findings that while Sardinians trace their ancestry mostly to the early Neolithic farmers, the French are more admixed with European hunter-gatherers and components of the Yamnaya expansion (Skoglund *et al.* 2014; Günther *et al.* 2015; Allentoft *et al.* 2015; Haak *et al.* 2015). To interpret the split times between Han, Dai and Karitiana, it should be noted that Karitiana is best modelled as a combination of three source populations (an ancient Siberian Eurasian source, a north East Asian source and an Australasian source), where the north East Asian contribution is substantially greater than the other two sources combined (Skoglund *et al.* 2015; Raghavan *et al.* 2015). The fact that the Karitiana show a more recent divergence with Han than with Dai likely reflects north East Asians contributing substantially to Native Americans, and that the Dai has a south East Asian component (closer to an Australasians, Skoglund *et al.* 2015; Raghavan *et al.* 2015). This admixture pattern results in shallower divergence between Karitiana and Han, and deeper divergence between Karitiana and Dai and between Han and Dai (see *e.g.* figure 2 in Raghavan *et al.* 2015).

### Western vs Eastern Africa and timing of the out-of-Africa event

Assuming a constant ancestral population size, we estimate the split between non-Africans and East-Africans (Dinka) to between 66 and 82 kya. The split between Mandenka and Yoruba is estimated to 100 kya while the split between Western Africans and Eastern Africans (Dinka and non-Africans) is estimated to between 96 and 117 kya.

The estimates based on the three outgroup ascertainment schemes 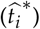 are generally older. Although the demographic history of Western and Eastern Africa appears to be particularly complex (Pickrell *et al.* 2014; Gurdasani *et al.* 2015; Triska *et al.* 2015; Busby *et al.* 2016; Hollfelder *et al.* 2017), and the standard deviation estimates suggest one should not over-interpret the 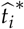 values, there are a few interesting tendencies among these estimates (figure 9). First, estimated split times are consistently lower among Yoruba, Mandenka and Dinka than between any of these populations and a non-African population; likely an effect of gene flow among the three African populations (Gurdasani *et al.* 2015; Busby *et al.* 2016; Schlebusch and Jakobsson 2018). Second, estimates between Yoruba and Dinka (or non-Africans) are deeper than split time estimates between Mandenka and Dinka (or non-Africans). This is consistent with some observations suggesting that Mandenka have a greater east African/European ancestry component compared to Yoruba (Gurdasani *et al.* 2015; Patin *et al.* 2017; Schlebusch and Jakobsson 2018). Although Mandenka is more distant geographically from Dinka than Yoruba, there is evidence that historical trading routes along the Sahel belt may have resulted in more gene-flow between East Africans and Mandenka (than with Yoruba) (Triska *et al.* 2015; Černý *et al.* 2018). Third, there is a tendency for split estimates between East Asians (Dai or Han) and West Africans (Yoruba, Mandenka or Dinka) to be deeper than split estimates between Europeans (French or Sardinian) and West Africans. This observation, combined with gene-flow between east and west Africa, is consistent with previous suggestions of migration into East Africa from a European or Middle Eastern source (Llorente *et al.* 2015).

**Figure 9.**
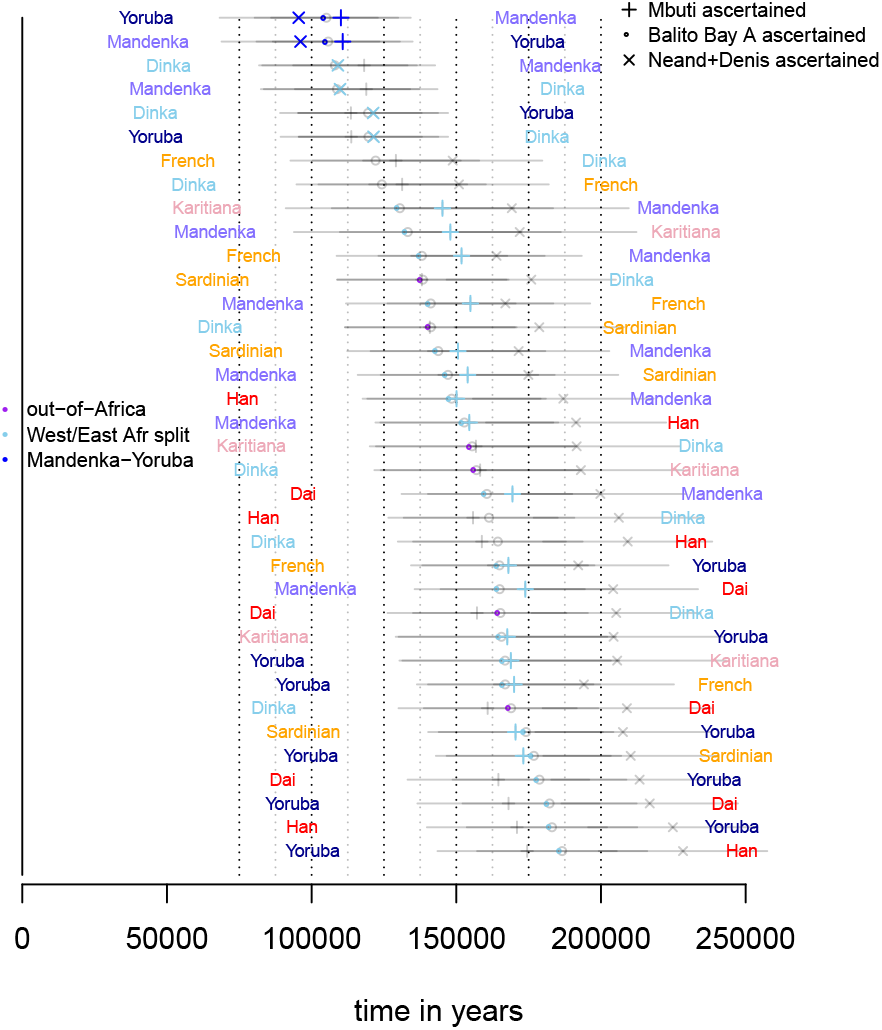
Different estimates of split times using outgroup ascertainment assuming a mutation rate of 1.25 × 10^−8^ and a generation time of 30 years. Three estimates are shown: estimates where outgroup ascertainment is performed in Mbuti (+), in Balito Bay A (o) and in Neandertal/Denisovan (×). Transparent grey represents standard deviation and for comparisons that failed the outgroup tests.

### Deepest splits among modern human populations

The split between Khoe-San and other modern human populations are estimated to around 250 kya using the TT-method (Schlebusch *et al.* 2017). It was further demonstrated that all modern-day Khoe-San groups and individuals, including the HGDP San individuals investigated here, were affected by Eurasian/east African admixture, which in turn impacts estimates of the deepest divergence of modern humans (different methods are differently sensitive to admixture), (Schlebusch *et al.* 2017). This observation became evident in comparisons with an ancient southern African individual (the Balito Bay A boy who lived some 2,000 years ago), closely related to modern-day Khoe-San individuals, but without the Eurasian/east African admixture that post-dated the life-time of the Balito Bay A boy. Population divergence time estimates based on the ancient Balito Bay A boy predates the estimates based on modern-day Khoe-San individuals, and give an estimate that is unaffected by the migration and admixture in the last 2000 years (Schlebusch *et al.* 2017).

Above we showed the effect of assuming a constant ancestral population size, and how violations of this assumption by a bottleneck in the ancestral population can bias divergence time estimates. However, we find no evidence for such a bottleneck in the common ancestral population to all modern humans, and hence

In general, we find very little difference between the TT and the TTo estimates (figure S19). In fact, there is a tendency for 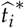 to be lower than 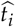 for the Mbuti-split, providing additional support that the Khoe-San split is the deepest split among modern human populations (Schlebusch *et al.* 2012).

### Archaic split times

The split between modern humans and both Neandertal and Denisovan is estimated to between 510 amd 707 kya, which is in line with previous such estimates (e.g., Prüfer *et al.* 2014). In fact, restricting the analysis to split times on the non-ancient branch (to alleviate issues with fossil dating and potential excess ancient DNA damage) and only to comparisons between Africans and the two archaic humans, gives a range of estimates from 609 to 681 kya (figure 10). Unfortunately, there is (presently) no suitable outgroup for comparisons between modern humans and archaic humans in order to utilize the outgroup ascertainment approach.

**Figure 10.**
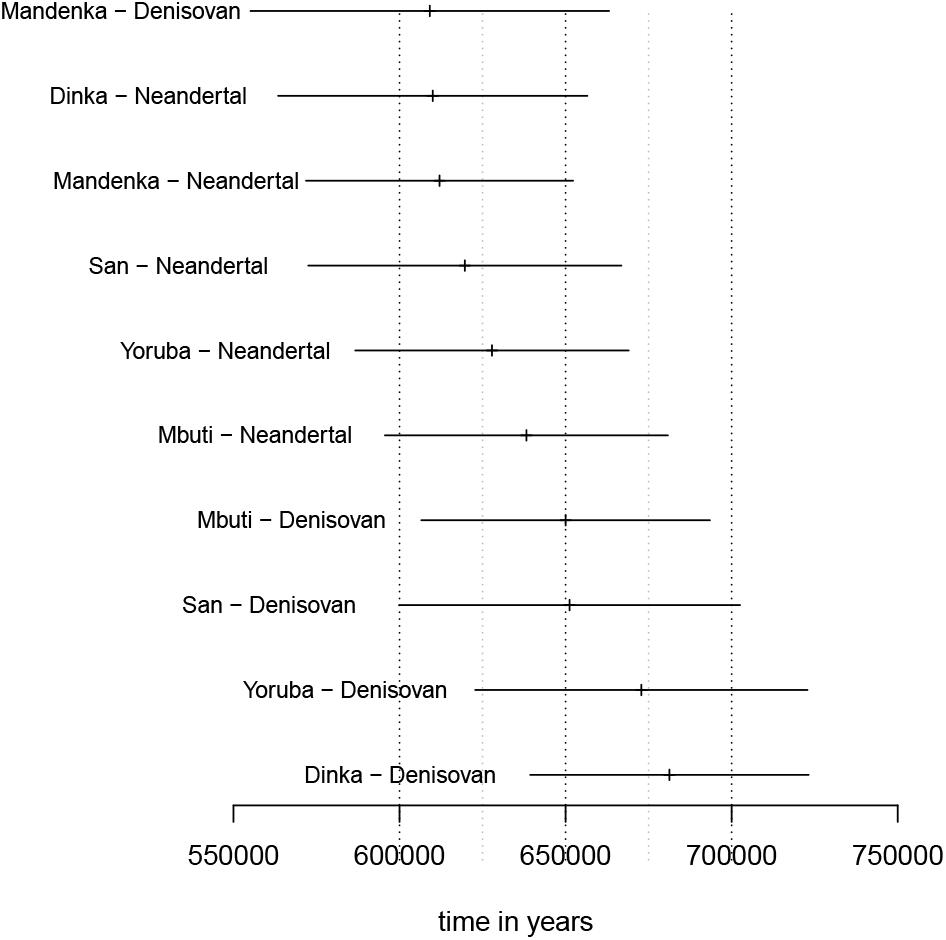
Split time estimates between the five African individuals and the two archaic humans assuming a constant ancestral population. Only estimates based on the non-ancient branch are shown. A mutation rate of 1.25 × 10^−8^ and a generation time of 30 years is assumed.

The estimated split between Neandertal and Denisovan is around 970 kya; more than 250 ky older than the split between modern humans and archaic humans. This is likely an artifact of violating the model assumptions; the existence of a more complex demography is indicated by our finding that neither Mbuti nor Balito Bay A were found to be true outgroups to the Neandertal-Denisovan comparison according to our outgroup test and the D-test (figures S14 and S13). Some studies hypothesize that the demographic relationship between Neandertals and Denisovans was governed by meta-population dynamics (Rogers *et al.* 2017). Others suggest complicating demographic factors such as admixture between Denisovans and *Homo erectus* (Prüfer *et al.* 2014) or admixture from the modern human branch into the Altai Neandertal (Kuhlwilm *et al.* 2016).

## Conclusion

We present a simple approach to estimate parameters under a comparatively general split model. In particular, no assumptions are needed concerning the population size processes/changes in the daughter populations (i.e., more recent than the split). The underlying model does not include gene-flow between daughter population, however, we can show that moderate violation of this assumption has little impact on the population divergence-time estimates. Assuming a constant ancestral population size, this approach provides an unbiased estimate of divergence time. However, when the ancestral population is not constant, and particularly in the case of severe bottlenecks, divergence time estimates can be biased. Indeed, simulations comparing the TT-method to GPhoCS – an alternative, fundamentally different approach to demographic inference, has shown that the two methods are sensitive to violations of the same assumptions. The reason for this can be intuitively understood in terms of the tMRCA in the ancestral population; most of this time is spent with two lineages and the duration of this is utilized by both methods to estimate the size of the ancestral population. Since, by assumption, the ancestral size is constant, if the time to the first coalescent event in the ancestral population is shorter than expected, (for instance, due to a bottleneck shortly before the divergence), then both methods underestimate the true population divergence time. When such severe bottlenecks have occurred, we have shown that it is possible to reduce much of this bias through the outgroup ascertainment procedure implemented in the TTo-method.

Applying the TT-method to a sample of 11 genomes from the HGDP panel together with the Neandertal and Denisovan genomes, we provide further information on the details of the various splits within the sample and corroborate many previously estimated population divergence times.

Finally, accumulating evidence suggests that human evolution is highly reticulated, and perhaps not well approximated by the sort of bifurcating tree-models studied here (Schlebusch and Jakobsson 2018; Henn *et al.* 2018; Scerri *et al.* 2018; Stringer 2016). Nonetheless, the framework presented here is still a useful tool: different population genetic methods vary in their assumptions and sensitivities to model violations, thus it is important when investigating the complex demographies underlying the evolution of humans to have access to a variety of different methods. The TT-method is relatively robust to model violations and provides a simple and transparent analytic framework that can be compared to, and potentially even integrated with, other, more computationally demanding methods (Beichman *et al.* 2017; Terhorst *et al.* 2017; Wang *et al.* 2020).

## Data availability

Genome sequence data from the following publications was extracted and reprocessed in order to avoid mapping and filtering biases: Prüfer *et al.* (2014); Schlebusch *et al.* (2017). Scripts used in simulations and plotting of results, together with open source code for running the TT method is freely available in Python at github.com/jammc313/TT-method.

## Appendix Ascertained data

Conditional on the derived variant being present in a true outgroup (in a population that has branched off before the investigated split), there are no mutations within the branches so that *μt*_1_, *μt*_2_, *μν*_1_ and *μν*_2_ can all be set to 0 in the equations 1 above. Thus we are left with

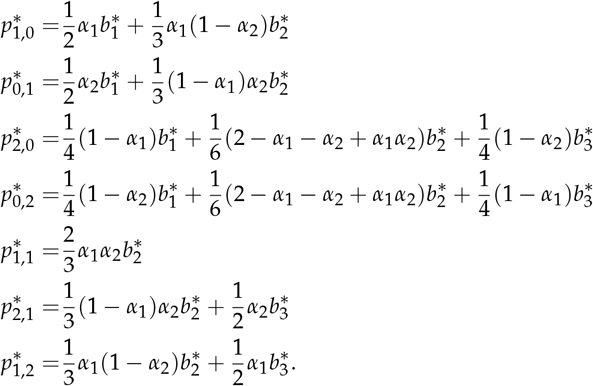

Under these equations,

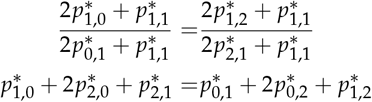

leading to the test statistics *Y*_1_ and *Y*_2_ in equations 5 and 6 above.

The second outgroup test (*Y*_2_) is similar to the D-test since the probability to draw the ancestral (single) allele from population 1 and the derived allele from population 2 is

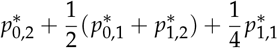

and the probability to draw the derived allele from population 1 and the ancestral allele from population 2 is

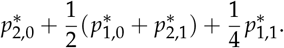

Similarly, for sites where the derived variant is observed in a sample of size 1 in the outgroup (note that our conditioning is different)

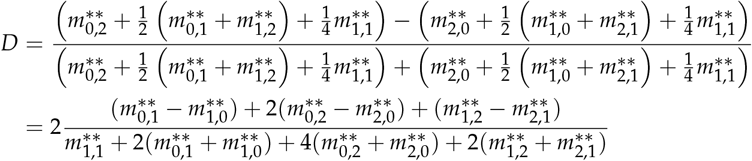

where ‘**’ indicates sites where the derived variant is observed in a sample of size 1 in the outgroup (our conditioning is different). The nominator is very similar to the nominator in the second outgroup test (*Y*_2_ in equation 6 above).

## Assuming an ancestral population with no structure backwards in time

Define

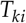

to be the number of generations a coalescent process that starts with *k* lineages at the (most recent) base of the ancestral population spends with *i* lineages (so that 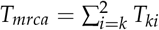). The probability that there are *k* derived variants in a sample of size *n* given that a mutation occurred when there were *i* lineages is (Slatkin 1996).

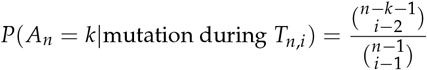

implying that

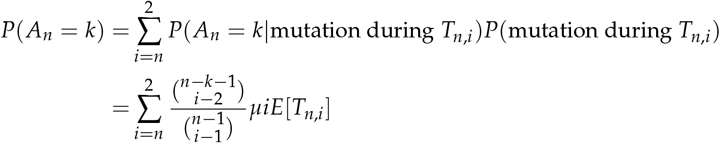

We get

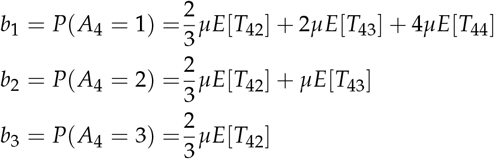

## Picking two gene copies from population 1 and one gene copy from population 2

Here, instead of two gene copies from both populations, the method assumes two sampled gene copies from population 1 and one sampled gene copy from population 2. The possible sample configurations are then:

**Table 3.** Number of derived in the two samples.

|  | 0 in pop2 | 1 in pop2 |
| --- | --- | --- |
| 0 in pop1 | $O_{0,0}$ | $O_{0,1}$ |
| 1 in pop1 | $O_{1,0}$ | $O_{1,1}$ |
| 2 in pop1 | $O_{2,0}$ | $O_{2,1}$ |

The observed number of sites with sample configuration *O*_*i,j*_ will be denoted by *m*_*i,j*_ and the total number of sites by *m*_*tot*_. Assume independence between sites, an infinite sites model and a split model with no migration where the two daughter populations merge into a panmictic ancestral population and define the event

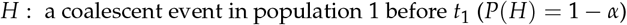

Also define *A*_*k*_ to be the number of derived variants in a (hypothetical) sample of size *k* drawn at the split time in the ancestral population. Writing *a*_*ki*_ = *P*(*A*_*k*_ = *i*), the conditional probabilities for sample configurations *O*_1,0_, · · ·, *O*_1,1_ are as in table 4:

**Table 4.** Conditional probabilities.

| | $H$ | $\neg H$ |
| --- | --- | --- |
| $O_{1,0}$ | $2\mu\nu$ | $\frac{2}{3}a_{31} + 2\mu t_1$ |
| $O_{0,1}$ | $\frac{1}{2}a_{21} + \mu t_2$ | $\frac{1}{3}a_{31} + \mu t_2$ |
| $O_{2,0}$ | $\frac{1}{2}a_{21} + \mu(t_1 - \nu)$ | $\frac{1}{3}a_{32}$ |
| $O_{1,1}$ | 0 | $\frac{2}{3}a_{32}$ |

Since

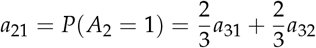

we write *b*_*i*_ = *a*_3*i*_ = *P*(*A*_3_ = *i*) to get

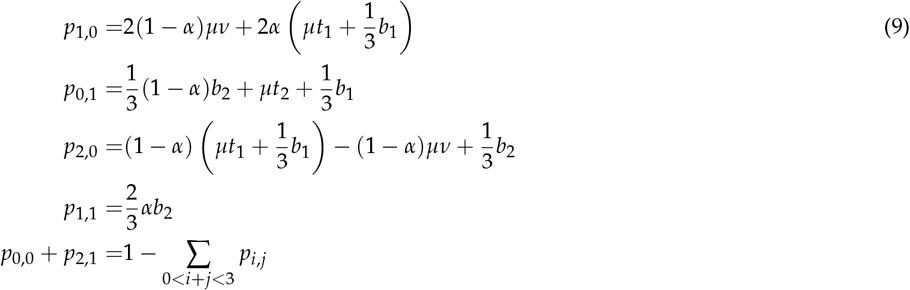

where *p*_*i,j*_ = *P*(*O*_*i,j*_).

Similar to the case when two gene copies are picked from each sub-population: 1) it is not possible to completely separate *b*_1_ from divergence times due to the co-occurrence of *b*_1_ with *μt*_1_ and *μt*_2_, 2) disregarding *b*_1_, this is an underdetermined set of equations with 5 parameters but only 4 equations/degrees of freedom (*p*_0,0_ + *p*_2,1_ = 1 − ∑_0<*i*+*j*<4_ *p*_*i,j*_). Setting *t*_1_ = *t*_2_ does not help since

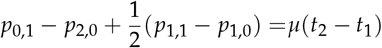

and thus reduces the number of independent equations. Assuming the model in figure 1b, does not reduce the number of parameters and does not help in this case.

If ascertainment is done in an outgroup

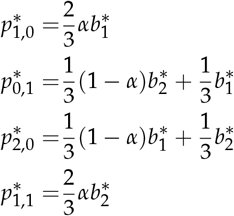

where ‘*’ indicates that these are the corresponding conditional parameters and probabilities. This can be solved to yield two estimates of *α*

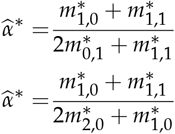

and these two estimates can be compared to create the tests

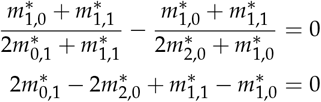

of that ascertainment was performed in a true outgroup.

Assuming the model in figure 1a, then *b*_2_ = *μE*[*T*_32_] and *b*_1_ = 3*μE*[*T*_33_] + *μE*[*T*_32_] and

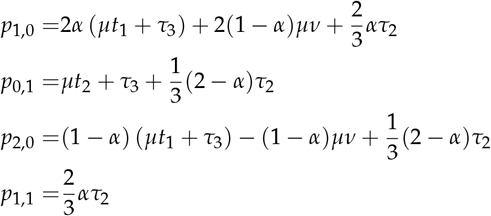

with *τ*_3_ = *μE*[*T*_33_] and *τ*_2_ = *μE*[*T*_32_]. If *α* is given by 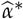, we solve to get

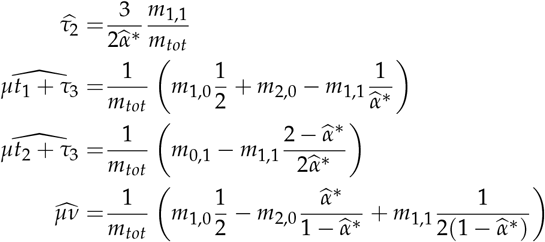

In (Schlebusch *et al.* 2012) and (Skoglund *et al.* 2011),

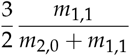

is used to estimate *α*. A comparison to the framework presented here gives

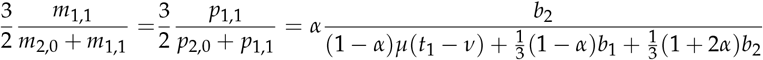

which is approximately *α* for *α* close to 1 or if *b*_1_ ≈ 2*b*_2_ and either *b*_2_ ≫ (1 − *α*)*μ*(*t*_1_ − *ν*) or *t*_1_ − *ν* ≈ 0.

Assuming a constant ancestral population size (figure 1b) implies *b*_1_ = 2*μN*_*A*_ = 2*b*_2_ and

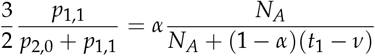

which is approximately *α* for *α* close to 1, for *N*_*A*_ ≫ (1 − *α*)(*t*_1_ − *ν*) and for *t*_1_ − *ν* ≈ 0.

If ascertainment is performed in an outgroup,

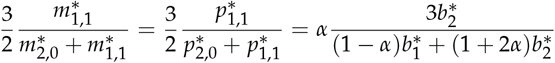

which is approximately *α* for *α* close to 1 or if 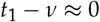

## Supplemental Material

### Figures for simulation studies

**Figure S1.**
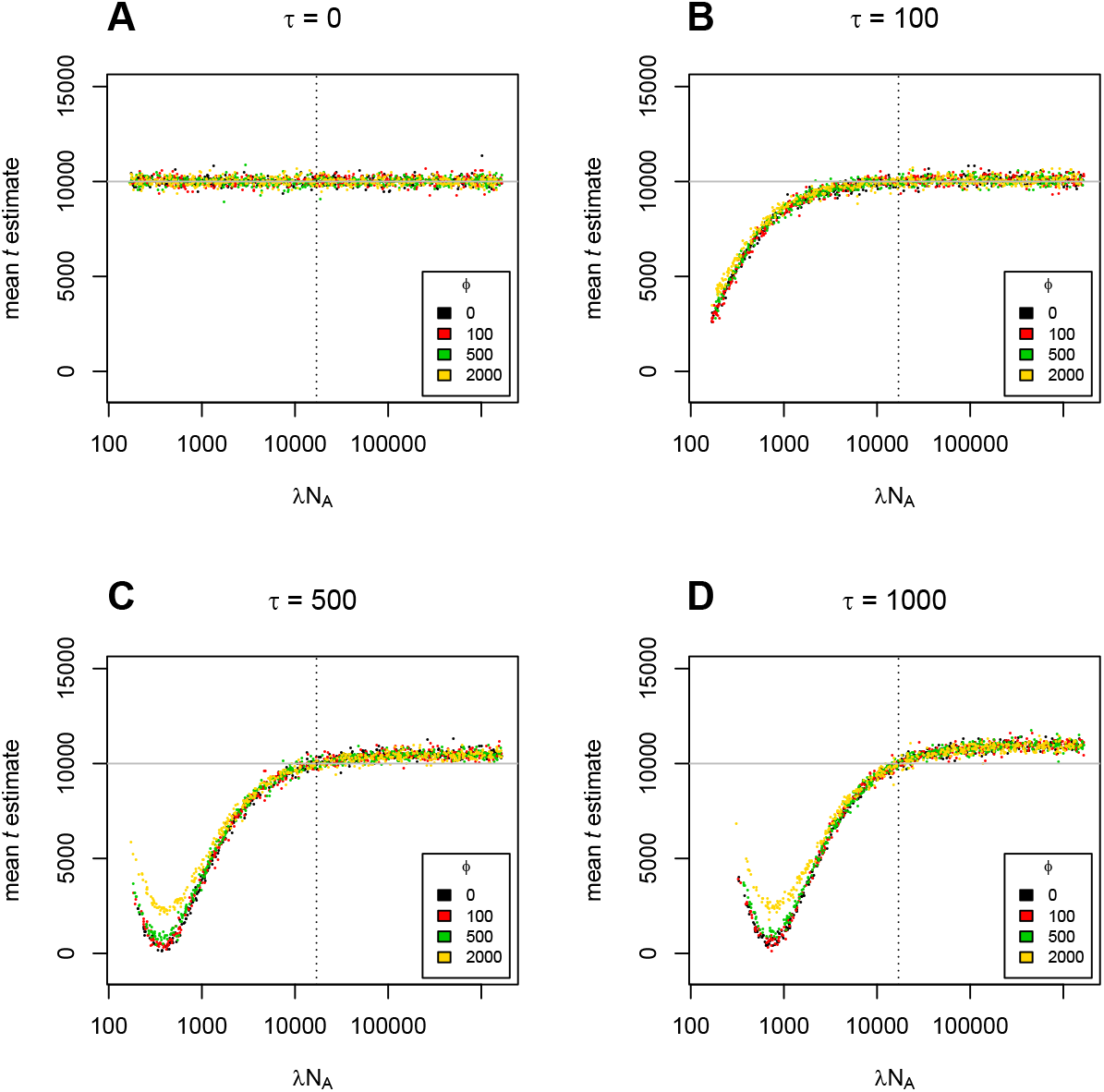
The effect of ancestral population size change, *λN*_*A*_, on TT method estimates of divergence time, *t*, with a true split time of 10,000 generations and ancestral population size change lasting (A) 0, (B) 100, (C) 500, and (D) 1000 generations.

**Figure S2.**
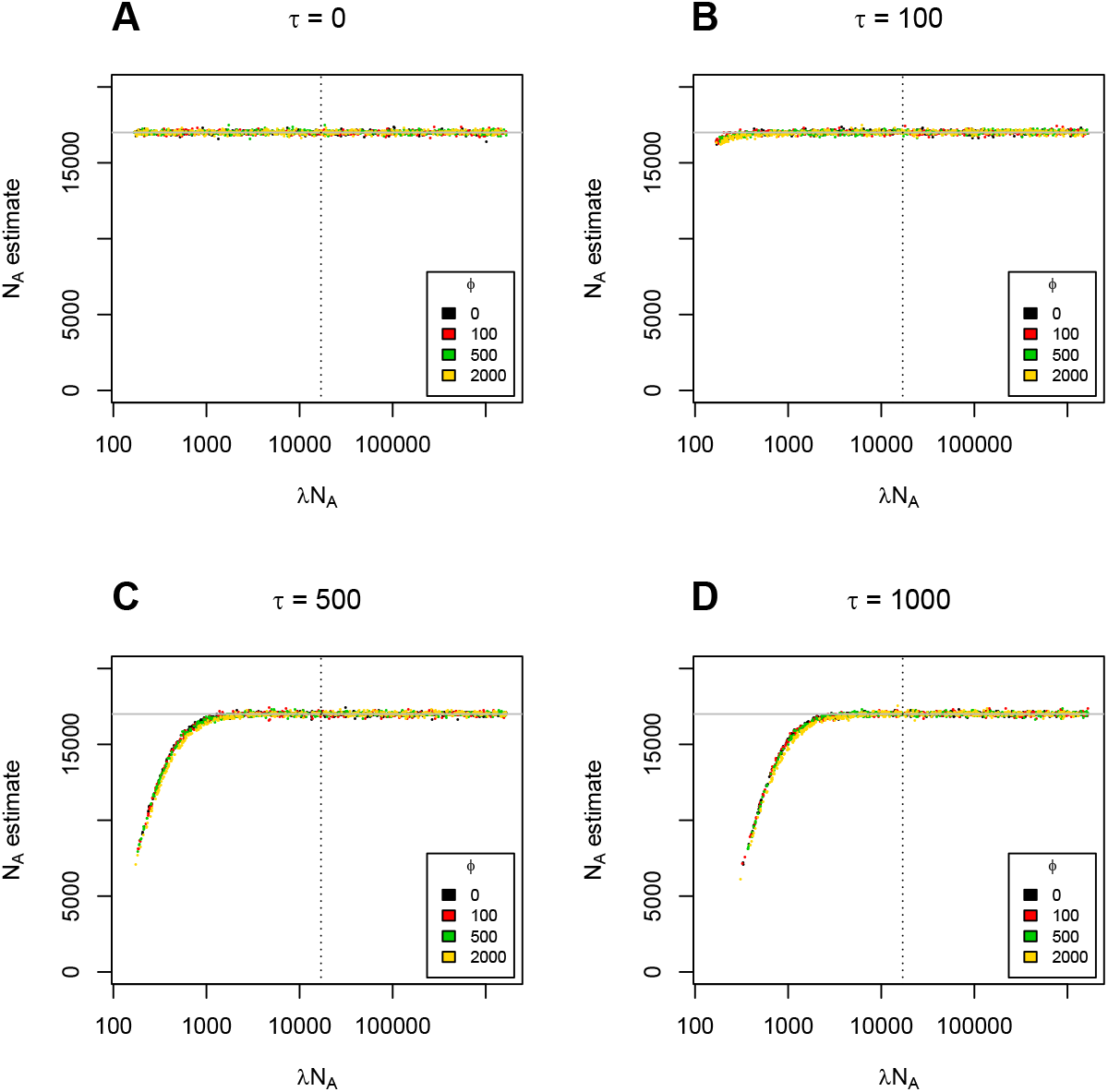
The effect of ancestral population size change, *λN*_*A*_, on TT method estimates of ancestral population size, *N*_*A*_, with a true split time of 10,000 generations and ancestral population size change lasting (A) 0, (B) 100, (C) 500, and (D) 1000 generations.

**Figure S3.**
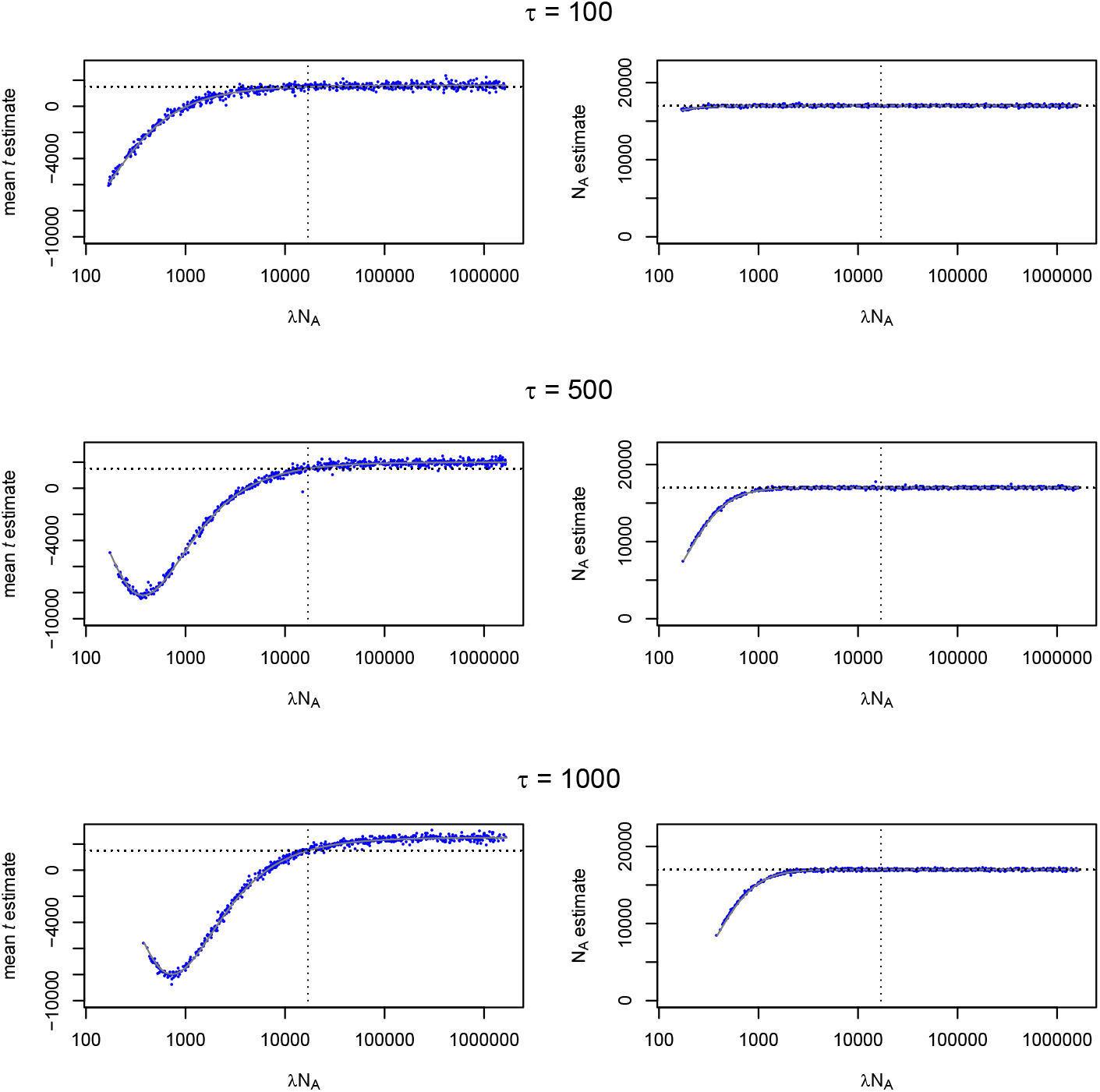
TT method estimates of divergence time and ancestral population size with changes to ancestral population size of duration (*τ*) (A, B) 100, (C, D) 500, and (E, F) 1,000 generations. In all cases, the change in ancestral population size occurs immediately prior to the split (*ϕ*=0) and the true split time is 1500 generations.

**Figure S4.**
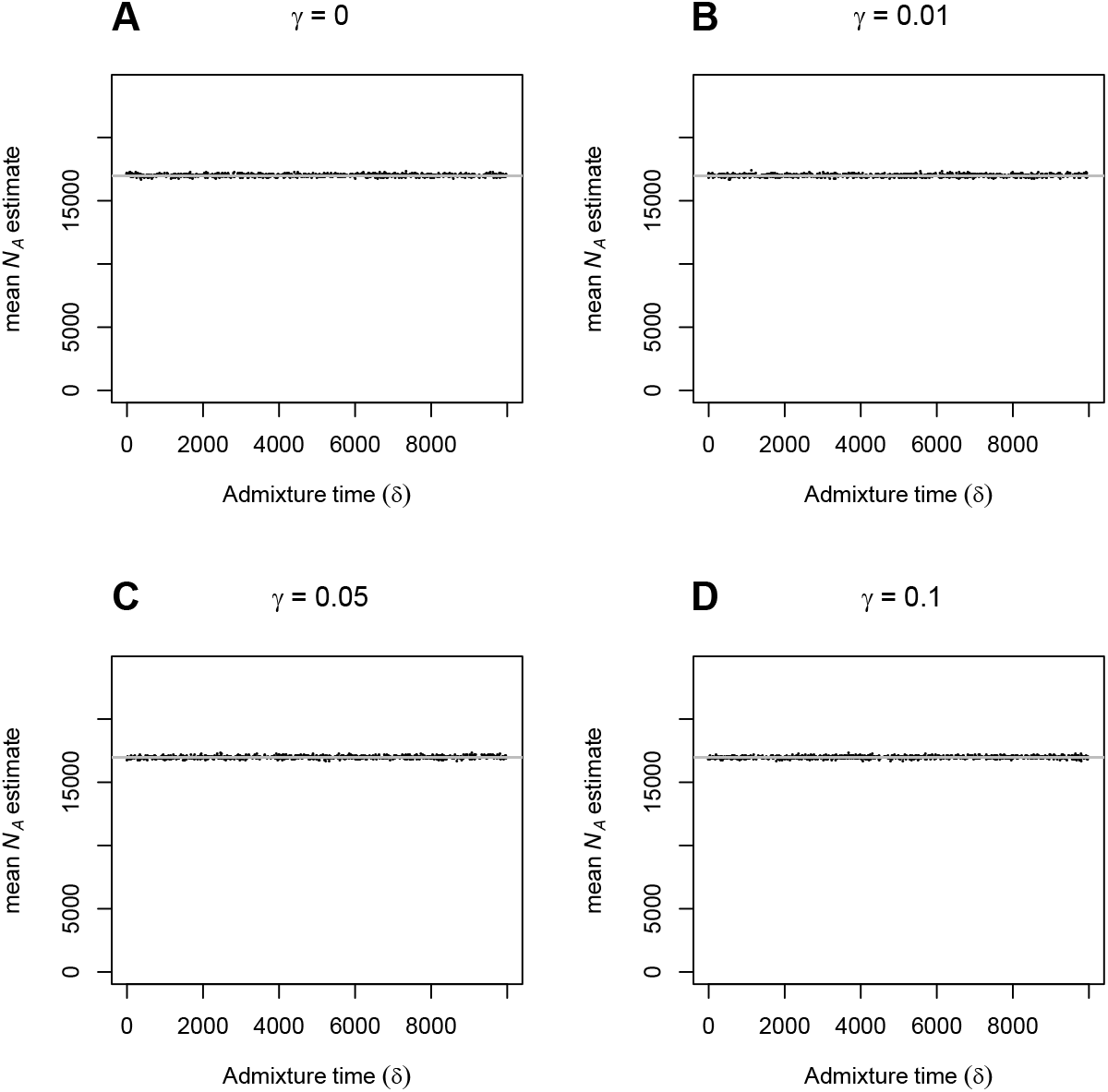
The effect of varying admixture time (*δ*) on TT method estimates of ancestral population size (*N*_*A*_), with migration proportions (*γ*) of (A) 0, (B) 0.01, (C) 0.05 and (D) 0.1 when true ancestral population size is 17,000 and true split time is 10,000 generations.

**Figure S5.**
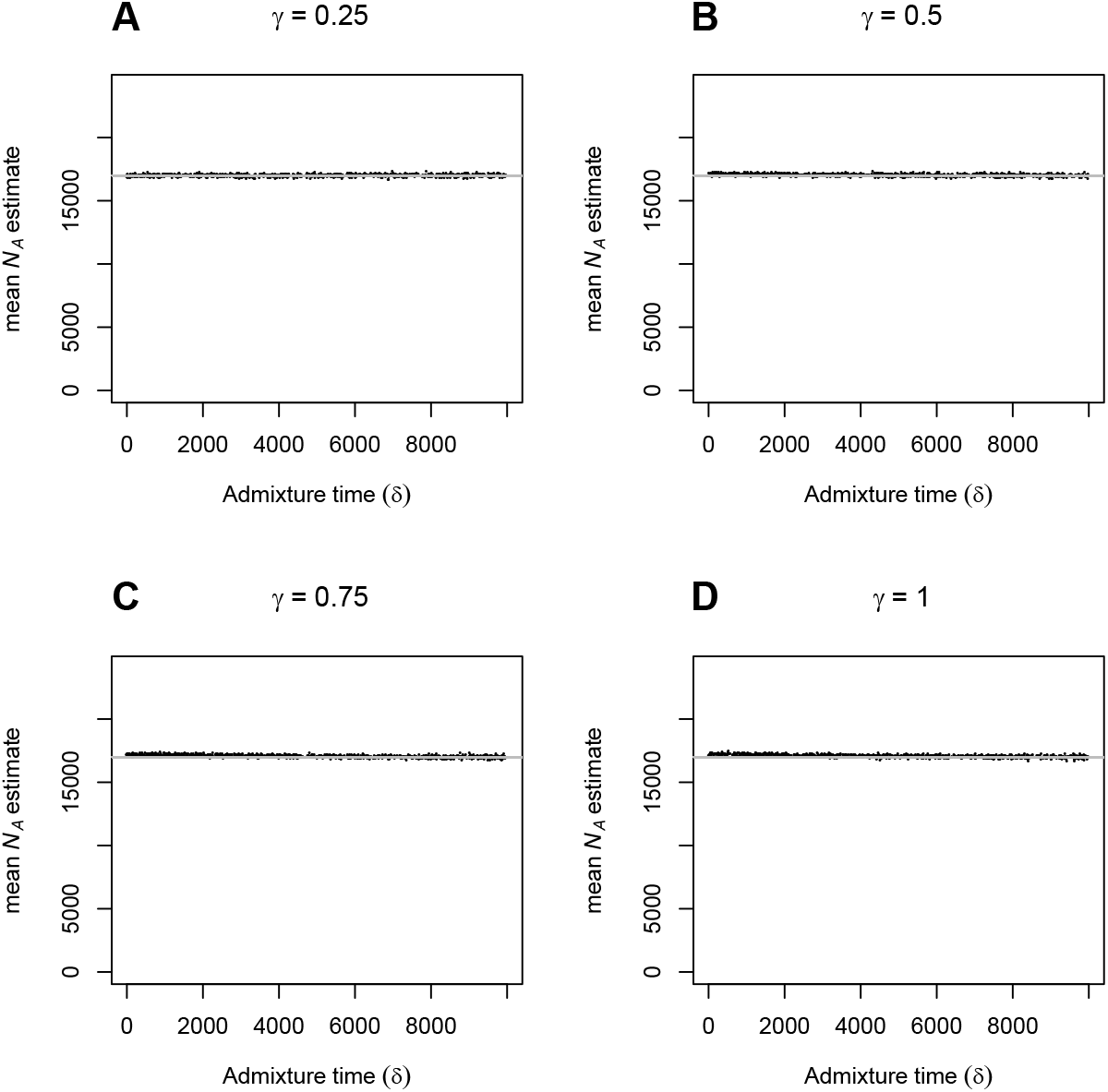
The effect of varying admixture time (*δ*) on TT method estimates of ancestral population size (*N*_*A*_), with migration proportions (*γ*) of (A) 0.25, (B) 0.5, (C) 0.75 and (D) 1 when true ancestral population size is 17,000 and true split time is 10,000 generations.

**Figure S6.**
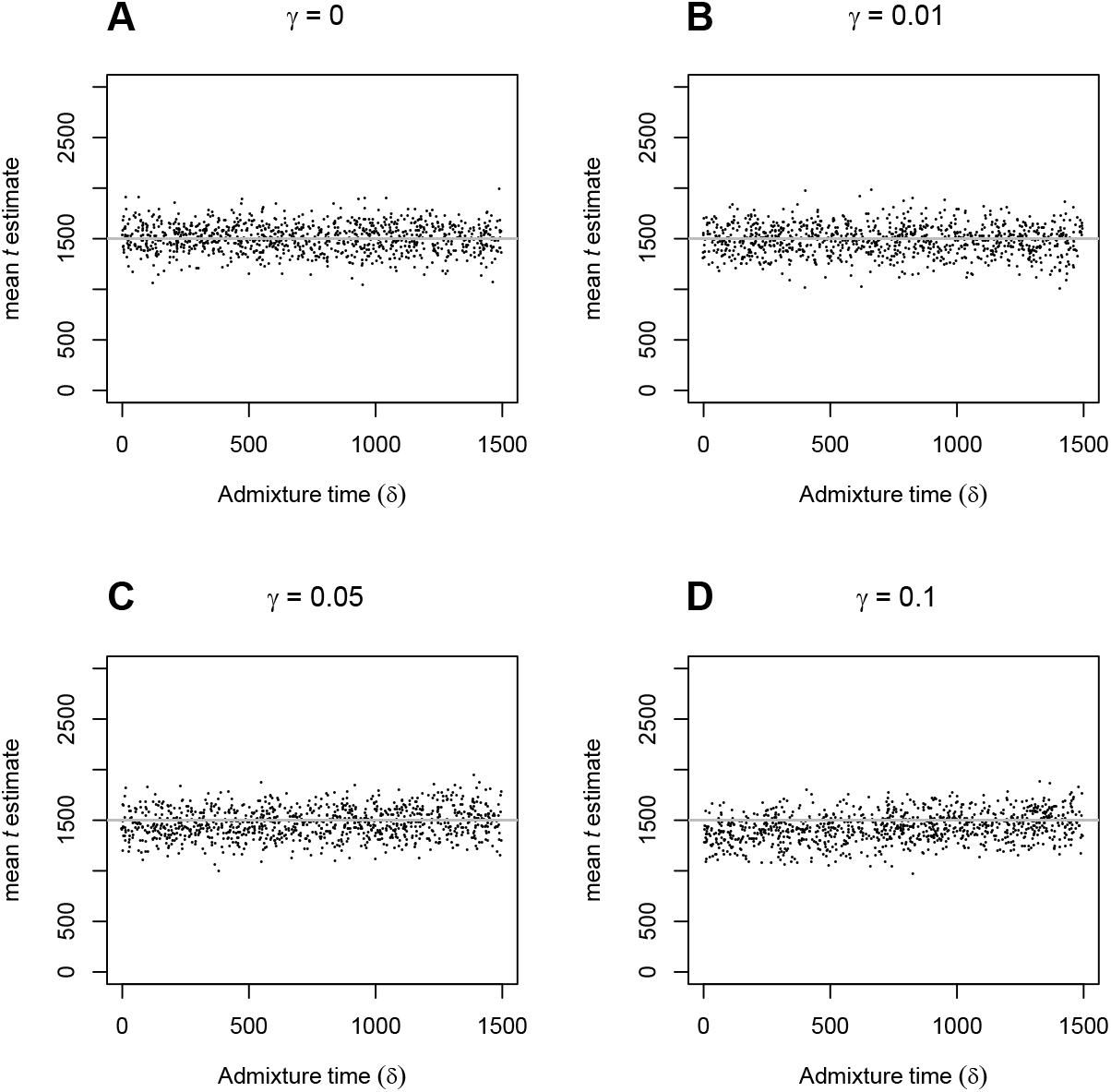
The effect of varying admixture time (*δ*) on TT split time estimates 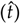, when the proportion of admixture (*γ*) is (A) 0, (B) 0.01, (C) 0.05 and (D) 0.1, and true split time is 1500 generations.

**Figure S7.**
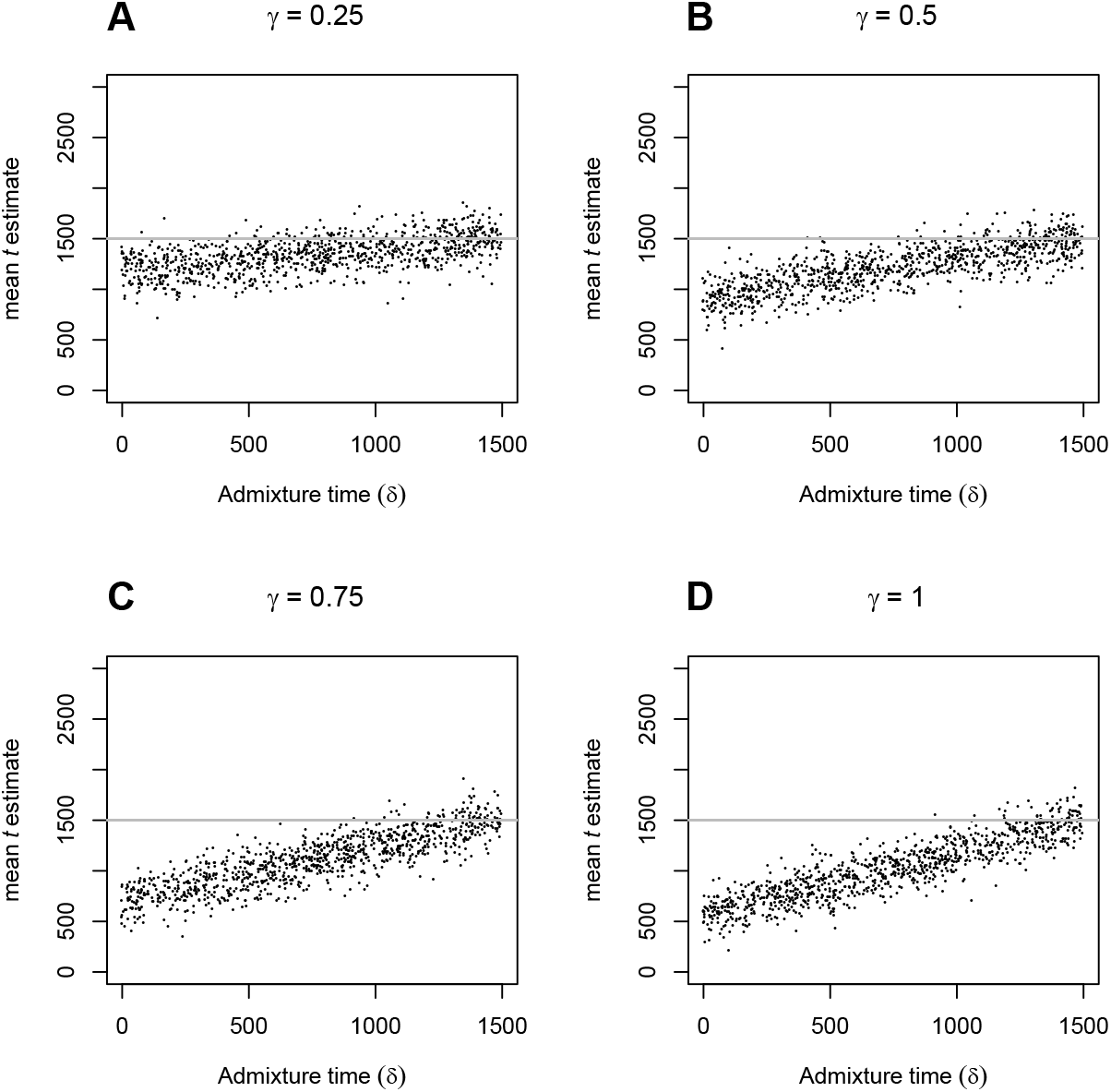
The effect of varying admixture time (*δ*) on TT split time estimates 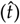, when the proportion of admixture (*γ*) is (A) 0.25, (B) 0.5, (C) 0.75 and (D) 1, and true split time is 1500 generations.

## Figures for application to data

### TT-estimates under a constant ancestral population

**Figure S8.**
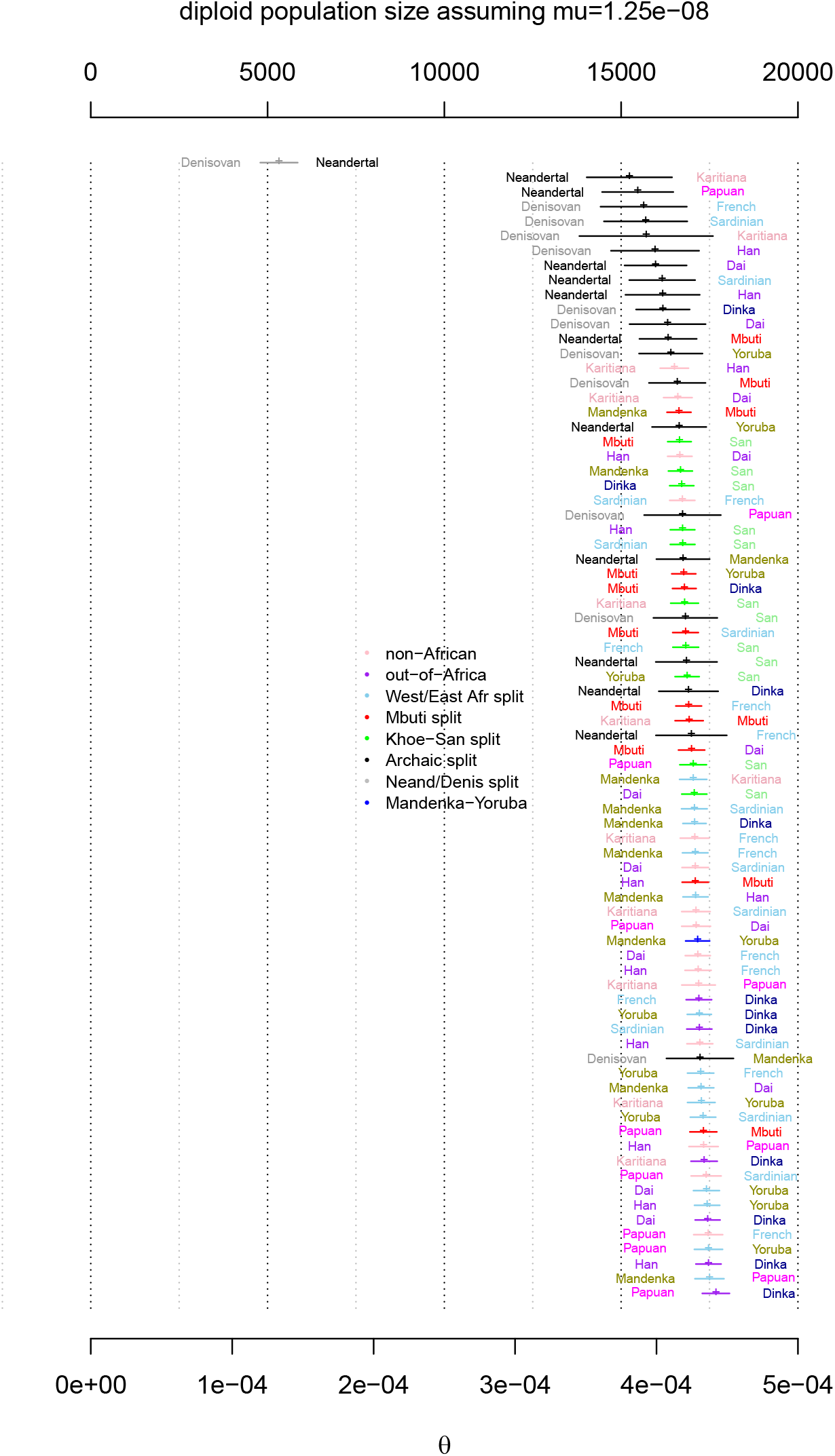
Estimates of *θ* = *μN*_*A*_ and the corresponding diploid ancestral population size (*N*_*A*_/2) assuming that it is constant and a mutation rate of 1.25 × 10^−8^.

**Figure S9.**
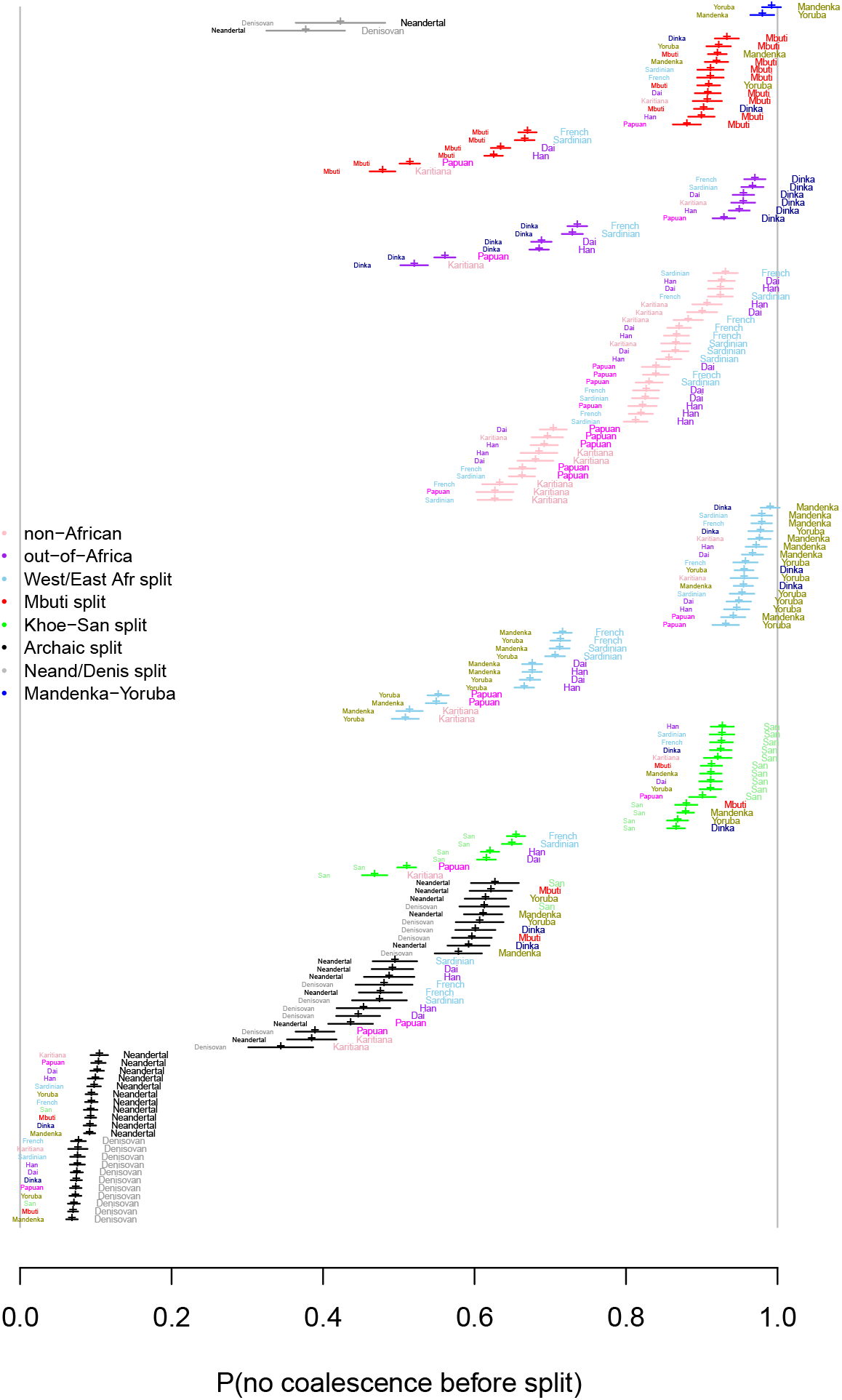
Estimates of *α* assuming a constant ancestral population.

**Figure S10.**
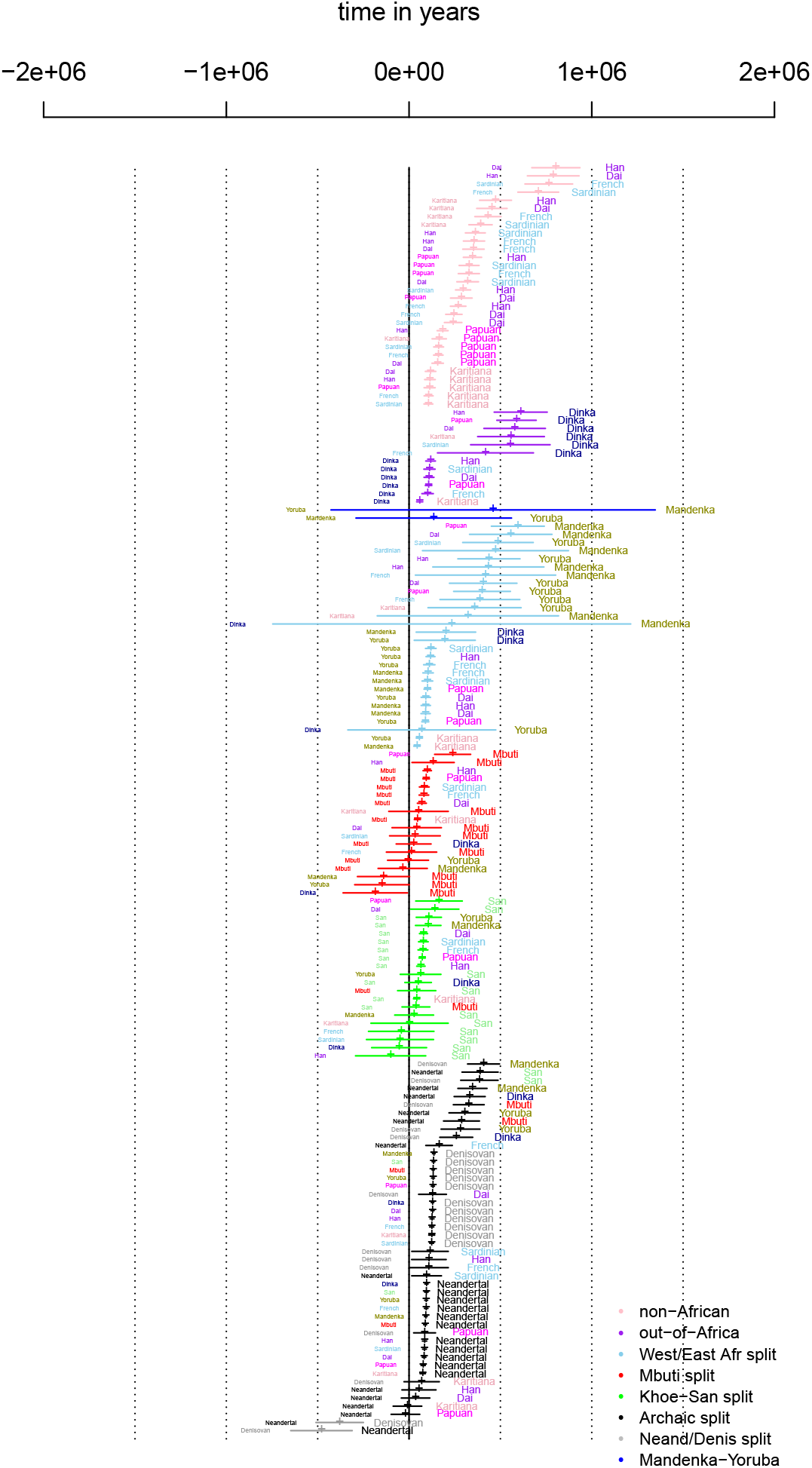
Estimates of expected coalescence times in years given coalescence before split assuming a constant ancestral population. A mutation rate of 1.25 × 10^−8^ and a generation time of 30 years is assumed.

**Figure S11.**
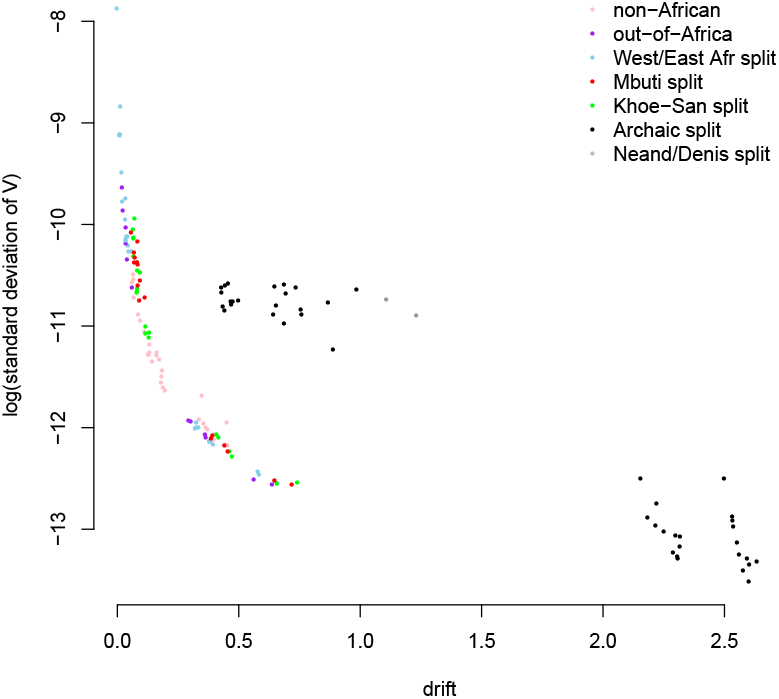
Drift (−*ln*(*α*)) vs estimated standard devation of *V* = *μν*.

**Figure S12.**
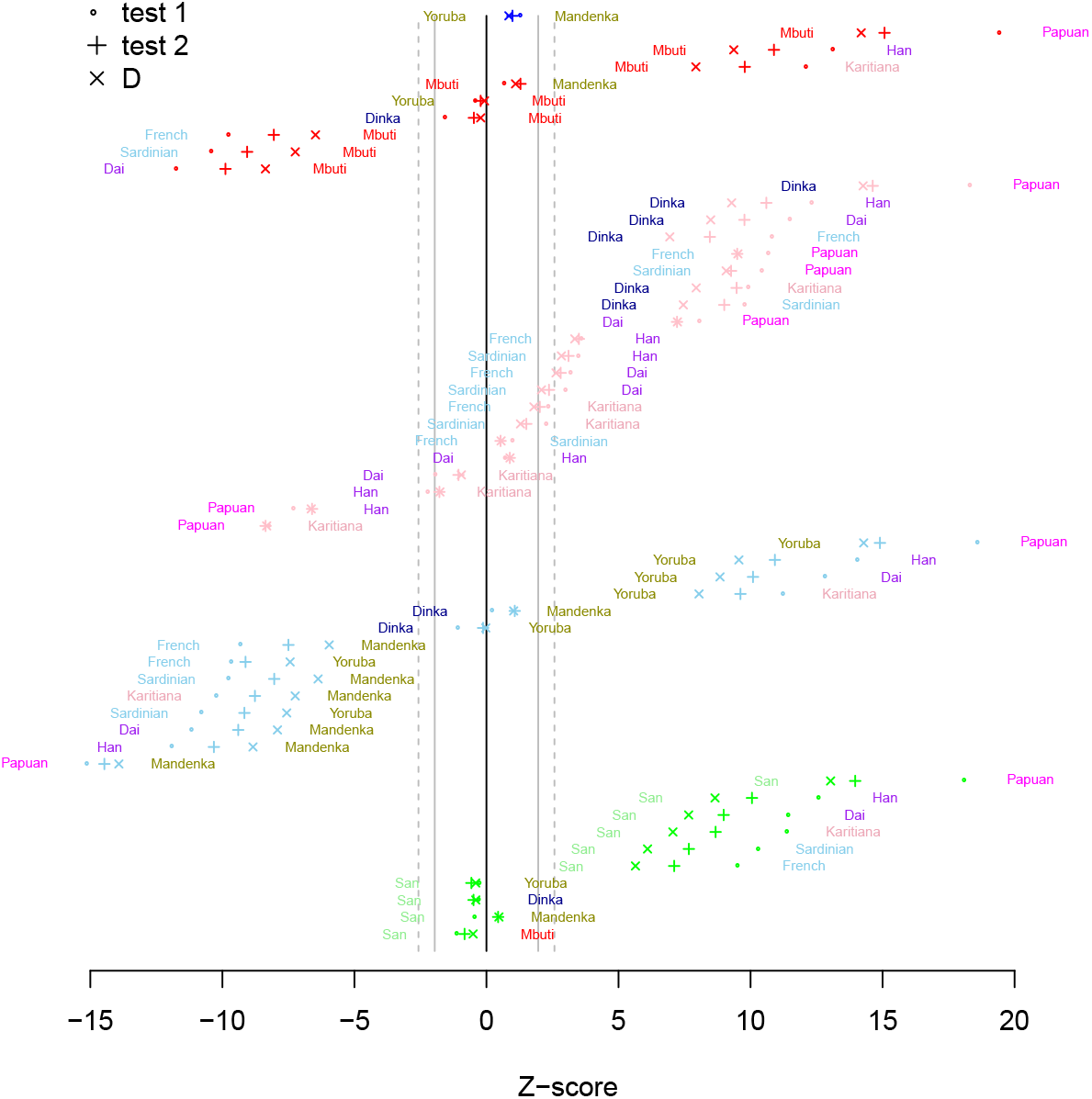
Tests based on equation 5 (○) and equation 6 (+) when data is conditional on the derived variant being present in either the Neandertal or Denisovan genome. D-tests (scaled by its estimated standard deviation, ×) with D(‘left population’, ‘right population‘, Neandertal+Denisovan, outgroup) are also shown. Solid grey lines shows cut-offs for rejecting null-model at the 0.05 level and the dashed lines at the 0.01 level.

**Figure S13.**
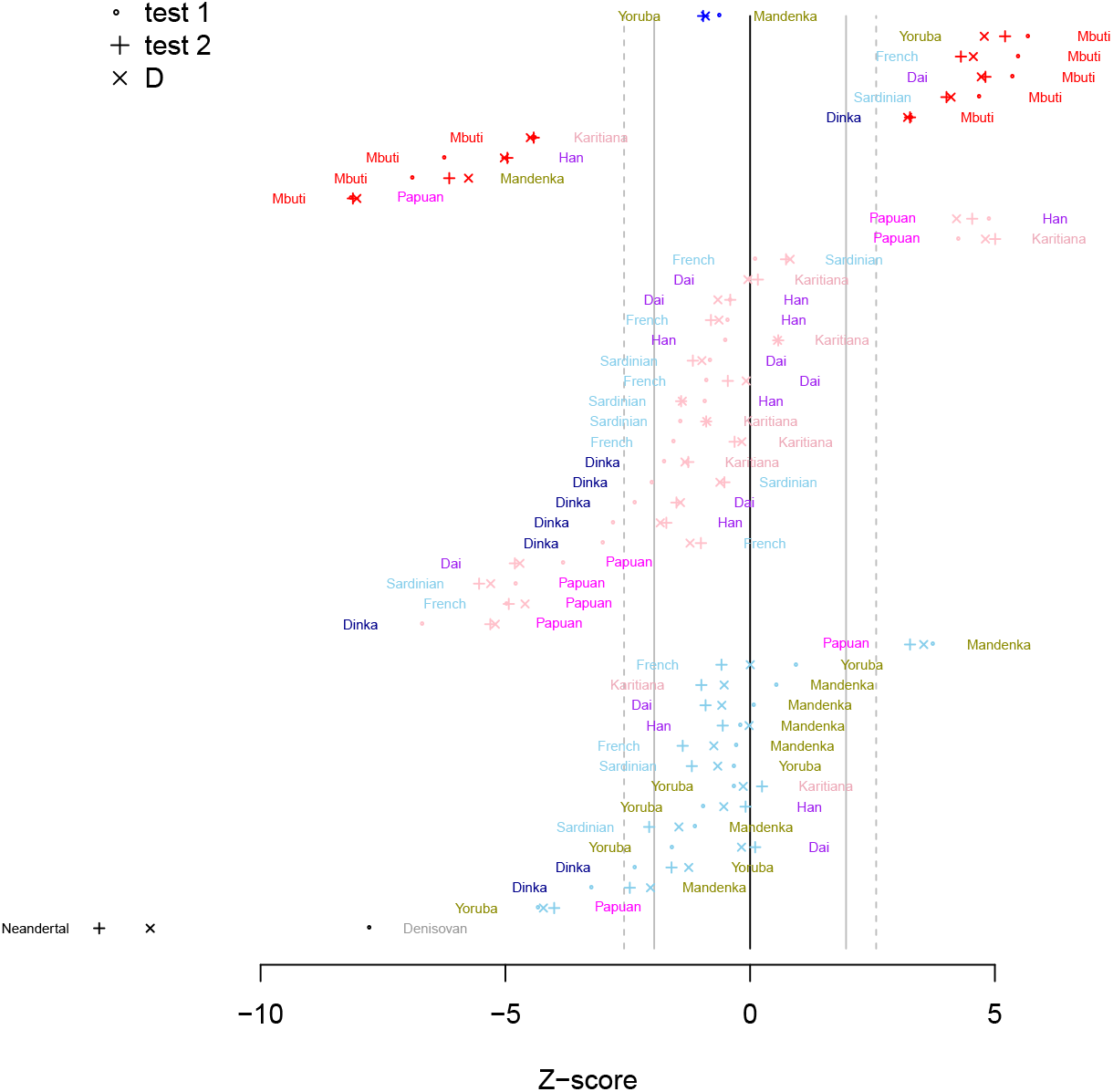
Tests based on equation 5 (○) and equation 6 (+) when data is conditional on the derived variant being present in Balito Bay A. D-tests (scaled by its estimated standard deviation, ×) with D(‘left population’, ‘right population’, Balito Bay A, outgroup) are also shown. Solid grey lines shows cut-offs for rejecting null-model at the 0.05 level and the dashed lines at the 0.01 level.

**Figure S14.**
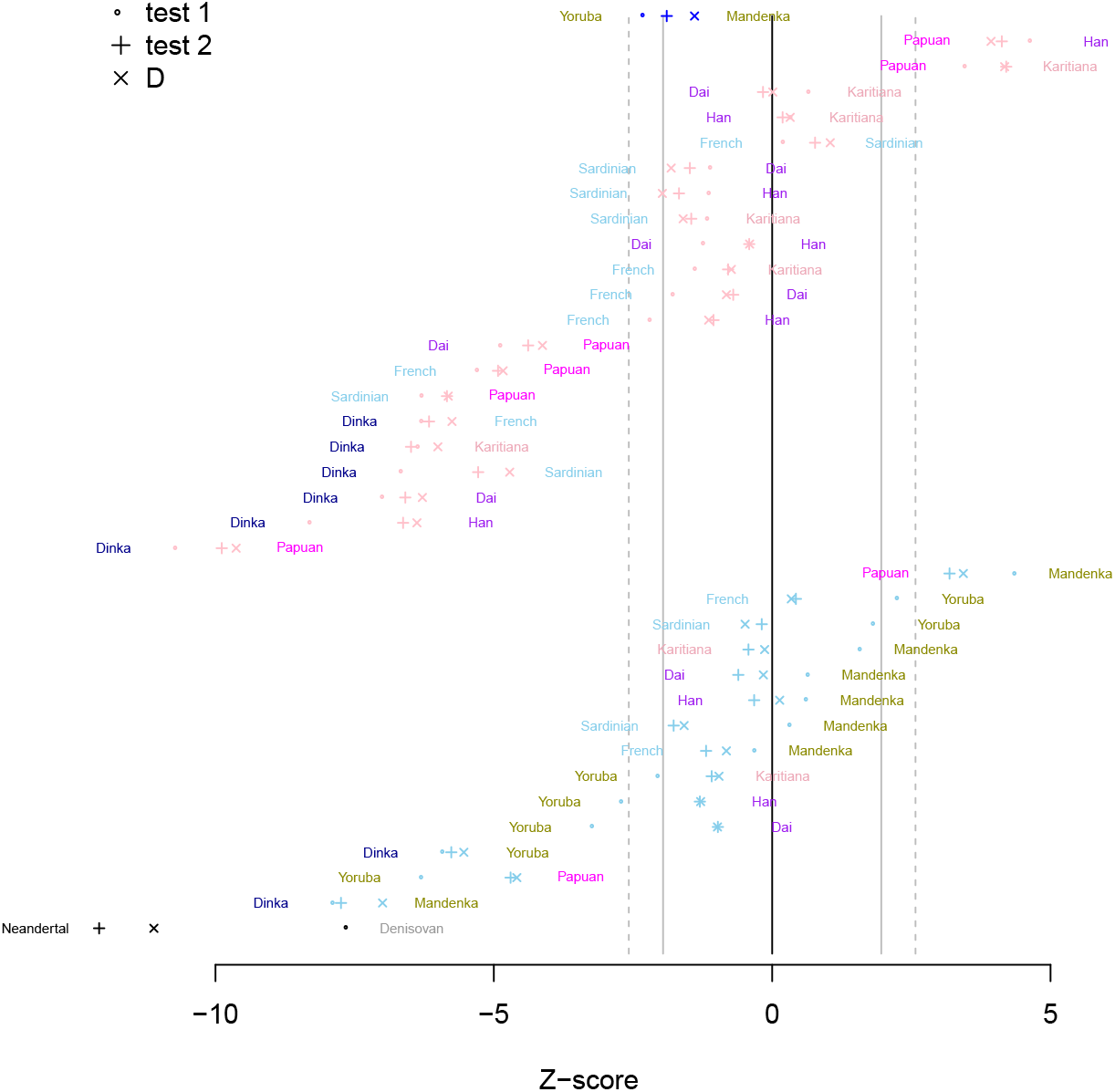
Tests based on equation 5 (○) and equation 6 (+) when data is conditional on the derived variant being present in Mbuti. D-tests (scaled by its estimated standard deviation, ×) with D(’left population’, ‘right population’, Mbuti, outgroup) are also shown. Solid grey lines shows cut-offs for rejecting null-model at the 0.05 level and the dashed lines at the 0.01 level.

**Figure S15.**
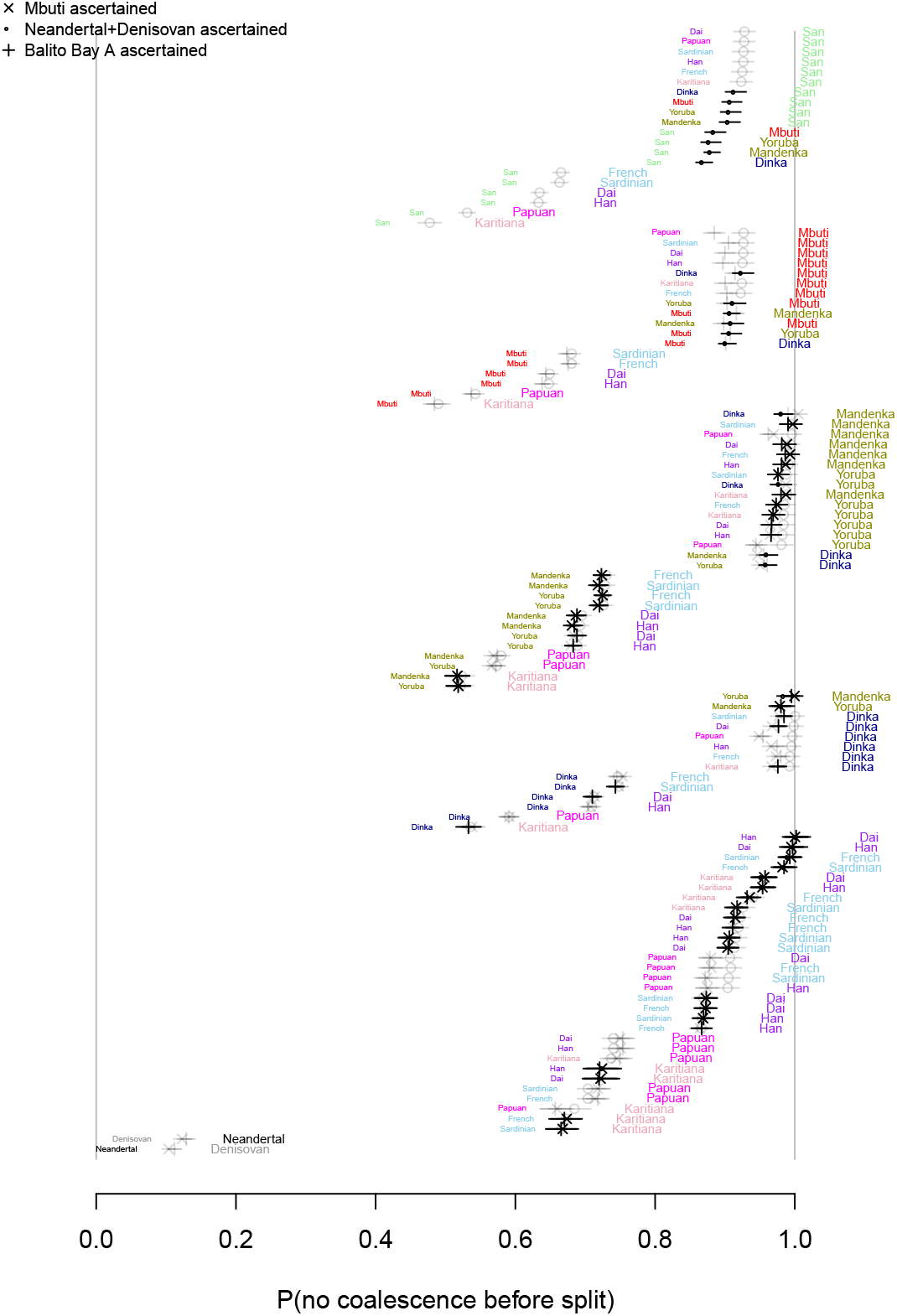
*α*-estimates conditional on the derived variant being present in the outgroup. ‘Cross’ show estimates when the derived variant is present in Mbuti, ‘circle’ show estimates when the derived variant is present in Neandertal or Denisovan, ‘plus’ show estimates when the derived variant is present in Balito Bay A. Black was used for estimates that did not fail the outgroup tests while transparent grey was used for comparisons that failed at least one of the two outgroup tests.

**Figure S16.**
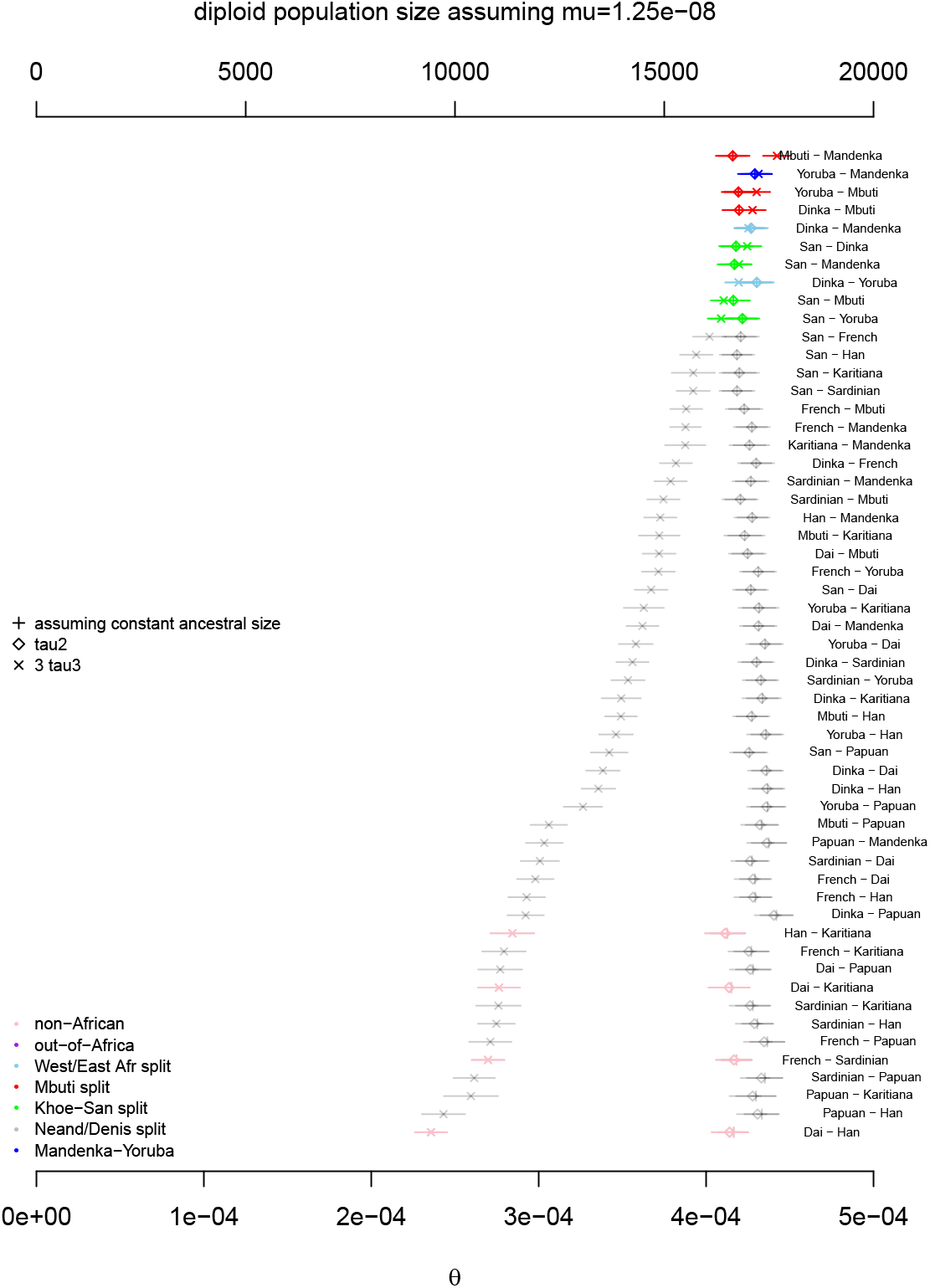
Different estimates of the ancestral population size. Here Neandertal+Denisovan was used as outgroup to estimate *τ*_2_ and *τ*_3_. Transparent grey was used for comparisons that failed at least one of the two outgroup tests.

**Figure S17.**
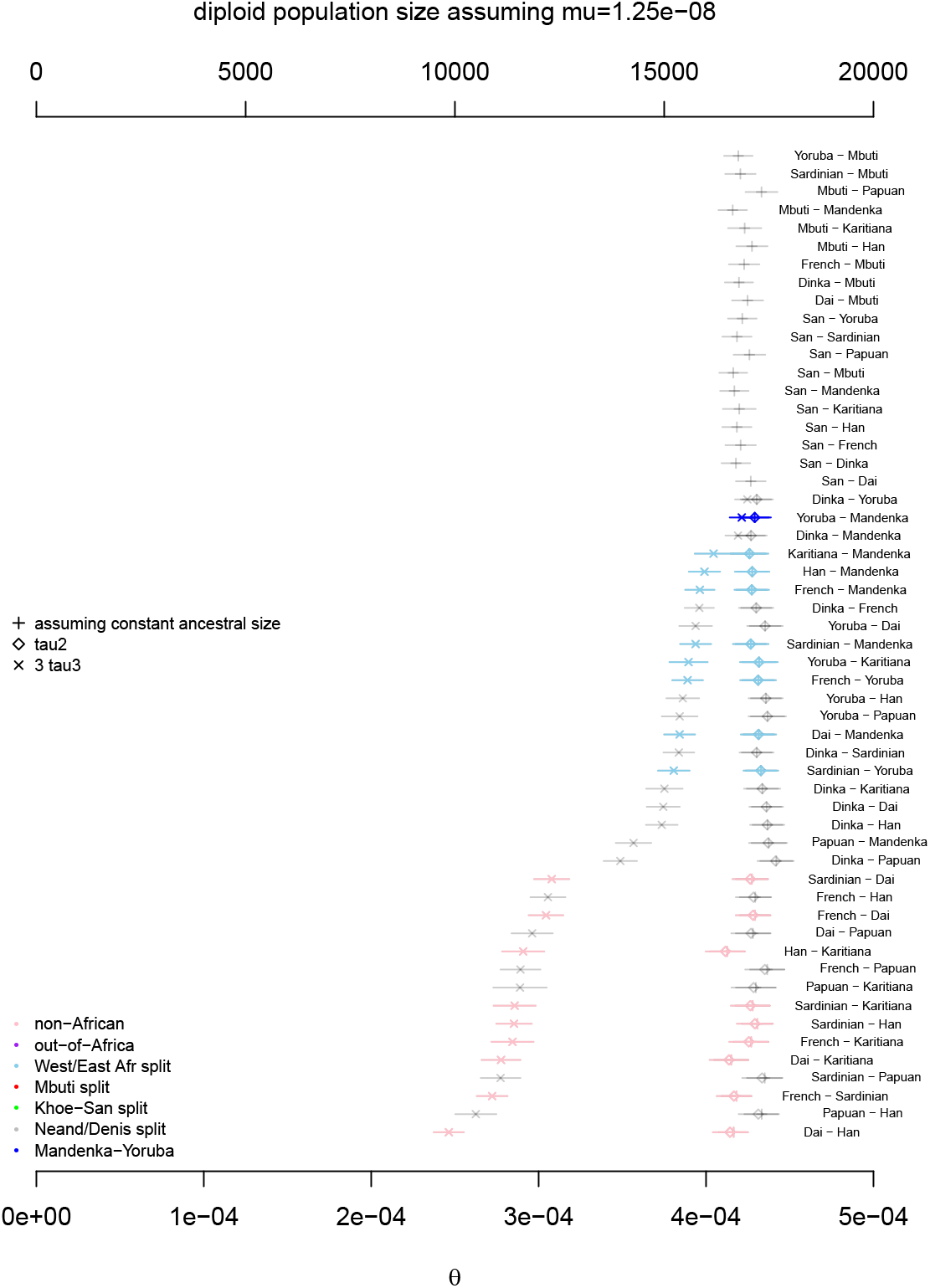
Different estimates of the ancestral population size. Here Mbuti was used as outgroup to estimate *τ*_2_ and *τ*_3_. Transparent grey was used for comparisons that failed at least one of the two outgroup tests.

**Figure S18.**
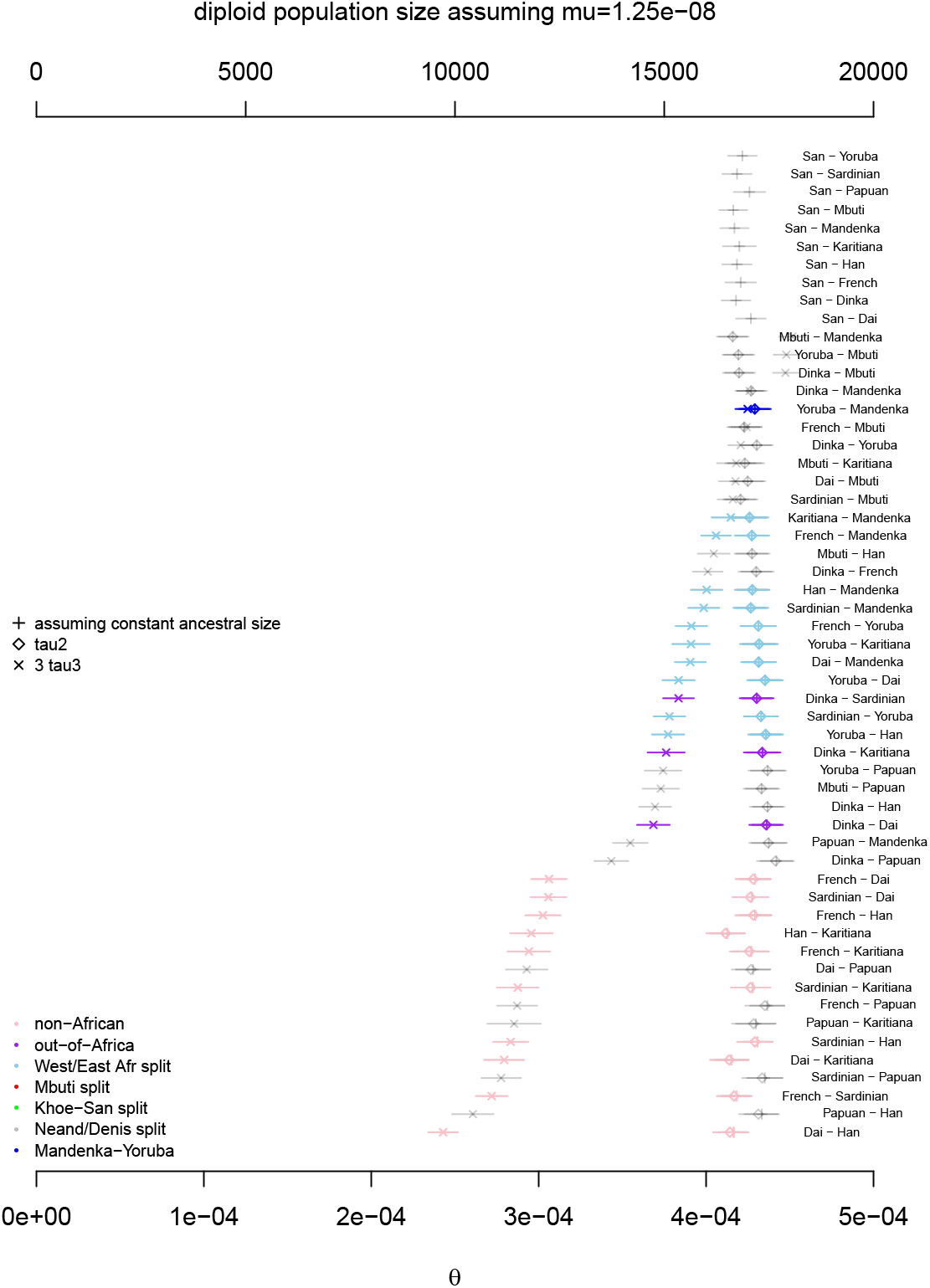
Different estimates of the ancestral population size. Here Balito Bay A was used as outgroup to estimate *τ*_2_ and *τ*_3_. Transparent grey was used for comparisons that failed at least one of the two outgroup tests.

**Figure S19.**
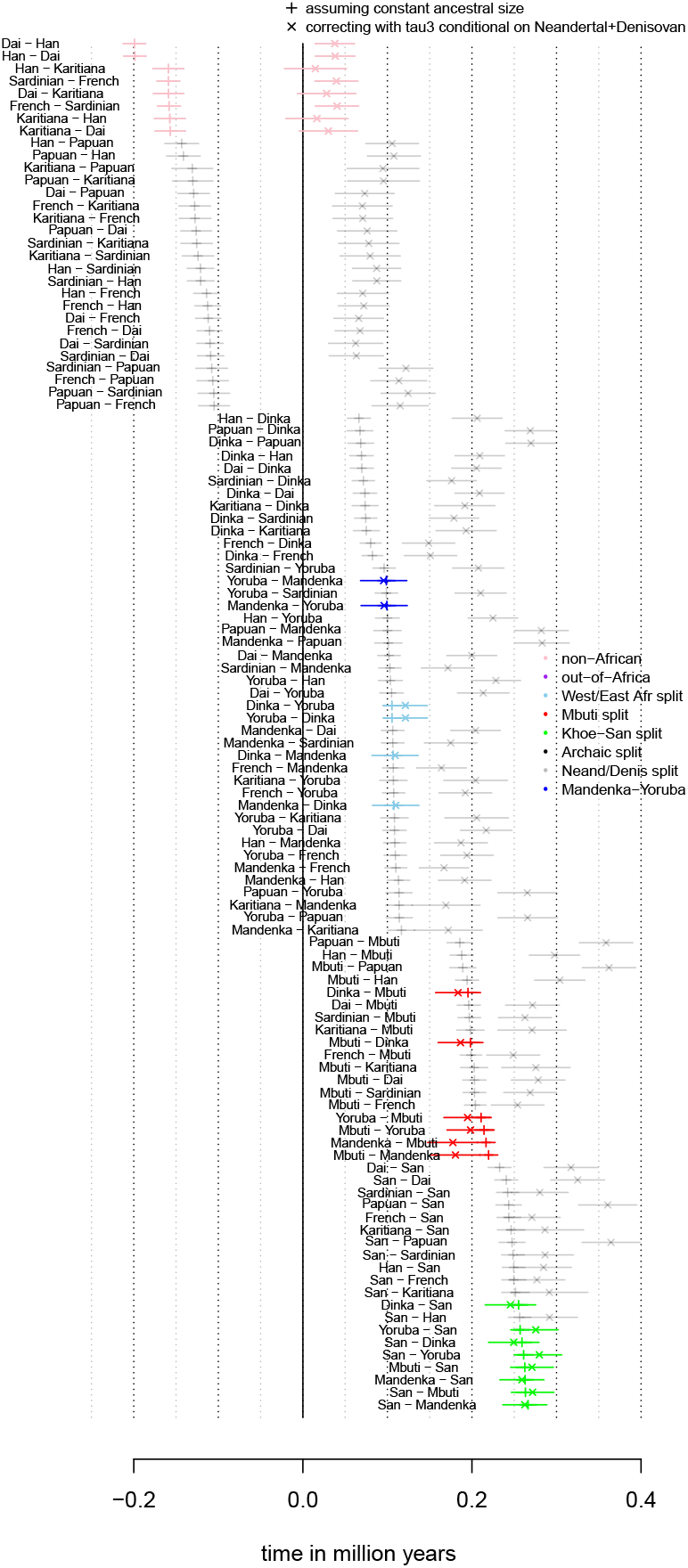
Different estimates of split times assuming a mutation rate of 1.25 × 10^−8^ and a generation time of 30 years. Two estimates are shown: estimates assuming a constant ancestral (+) and estimates relying on an external estimate of *α* (×) obtained by outgroup ascertainment in Neandertal+Denisovan. Transparent grey was used for comparisons that failed at least one of the two outgroup tests.

**Figure S20.**
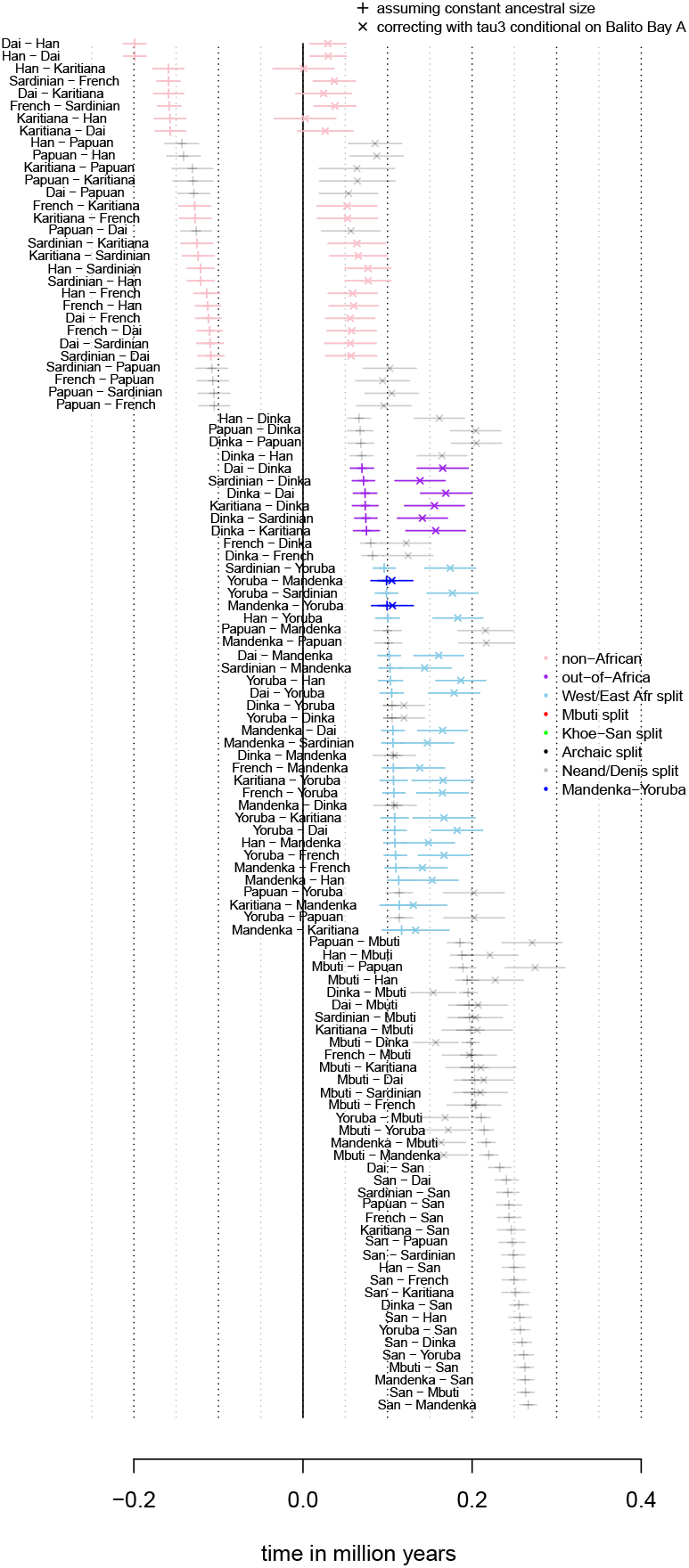
Different estimates of split times assuming a mutation rate of 1.25 × 10^−8^ and a generation time of 30 years. Two estimates are shown: estimates assuming a constant ancestral (+) and estimates relying on an external estimate of *α* (×) obtained by outgroup ascertainment in Balito Bay A (an ancient Khoe-San genome). Transparent grey was used for comparisons that failed at least one of the two outgroup tests.

**Figure S21.**
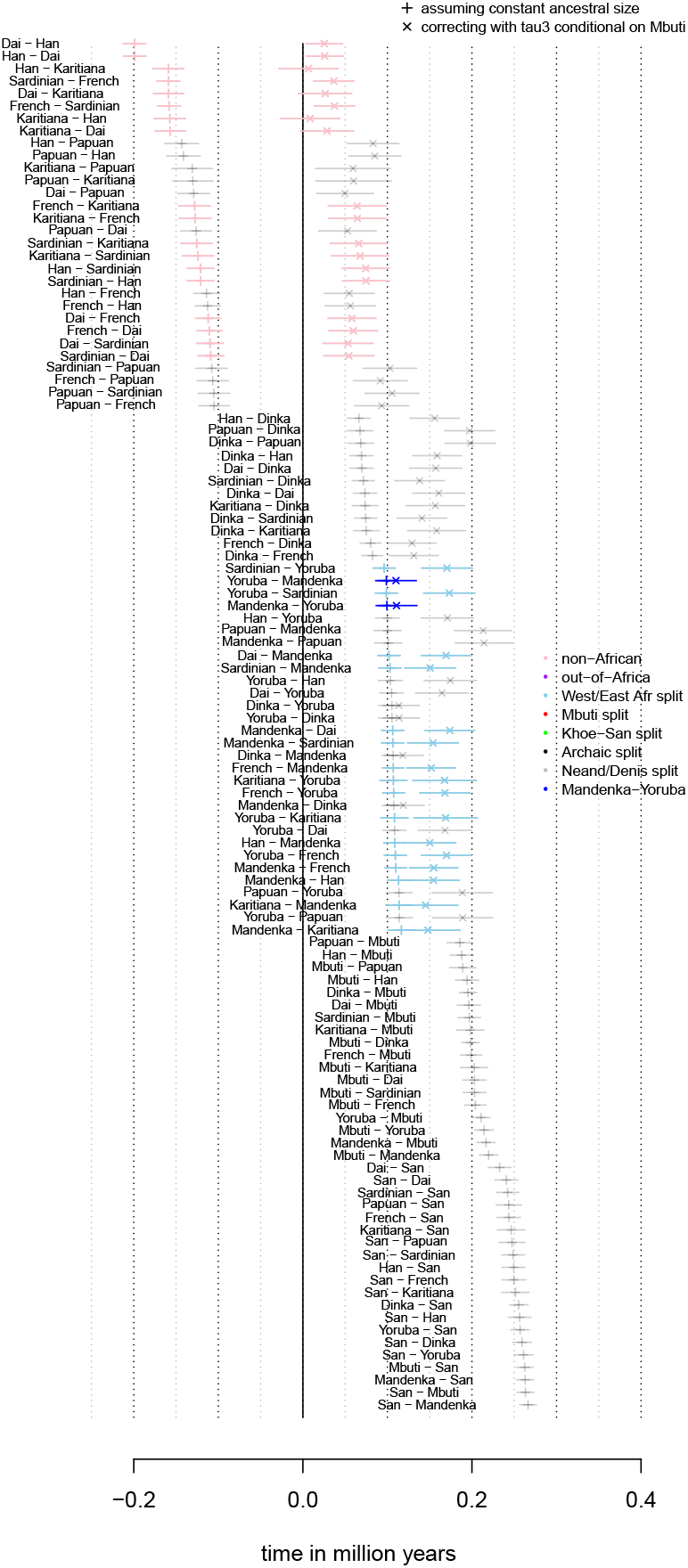
Different estimates of split times assuming a mutation rate of 1.25 × 10^−8^ and a generation time of 30 years. Two estimates are shown: estimates assuming a constant ancestral (+) and estimates relying on an external estimate of *α* (×) obtained by outgroup ascertainment in Mbuti. Transparent grey was used for comparisons that failed at least one of the two outgroup tests.

## Literature Cited

Allentoft, M. E., M. Sikora, K.-G. Sjögren, S. Rasmussen, M. Ras-mussen, et al., 2015 Population genomics of bronze age eurasia. Nature 522: 167.

Beaumont, M., A. Zhang, and W. Balding, 2002 Approximate Bayesian computation in population genetics. Genetics 162: 2025–2035.

Beichman, A. C., T. N. Phung, and K. E. Lohmueller, 2017 Com-parison of single genome and allele frequency data reveals discordant demographic histories. G3: Genes, Genomes, Genetics 7: 3605–3620.

Busby, G. B., G. Band, Q. Si Le, M. Jallow, E. Bougama, et al., 2016 Admixture into and within sub-saharan africa. eLife 5: e15266.

Busing, F. M. T. A., E. Meijer, and R. V. D. Leeden, 1999 Delete-m jackknife for unequal m. Statistics and Computing 9: 3–8.

Chen, H., 2012 The joint allele frequency spectrum of multi-ple populations: A coalescent theory approach. Theoretical Population Biology 81: 179–195.

Cornuet, J., M. Pudlo, P. Veyssier, J. Dehne-Garcia, A. Gautier, et al., 2014 DIYABC v2.0: a software to make approx-imate Bayesian computation inferences about population history using single nucleotide polymorphism, DNA se-quence and microsatellite data. Bioinformatics 30: 1187–1189, doi:10.1093/bioinformatics/btt763.

Doob, J. L., 1934 Probability and statistics. Transactions of the American Mathematical Society 36: 759–775.

Excoffier, L., I. Dupanloup, E. Huerta-Sánchez, V. C. Sousa, and M. Foll, 2013 Robust demographic inference from genomic and SNP data. PLoS Genet. 9: e1003905.

Gattepaille, L., T. Günther, and M. Jakobsson, 2016 Inferring past effective population size from distributions of coalescent times. Genetics 204: 1191–1206.

Green, R. E., J. Krause, A. W. Briggs, T. Maricic, U. Stenzel, et al., 2010 A Draft Sequence of the Neandertal Genome. Science 328: 710–722.

Griffiths, R. and S. Tavaré, 1998 The age of a mutation in a general coalescent tree. Communications in Statistics. Stochastic Models 14: 273–295.

Gronau, I., M. Hubisz, J. Gulko, B. Danko, and C. Siepel, 2011 Bayesian inference of ancient human demography from individual genome sequences. Nature Genetics 43: 1031–1035, doi:10.1038/ng.937.

Günther, T., C. Valdiosera, H. Malmström, I. Ureña, R. Rodriguez-Varela, et al., 2015 Ancient genomes link early farmers from atapuerca in spain to modern-day basques. Proceedings of the National Academy of Sciences 112: 11917–11922.

Gurdasani, D., T. Carstensen, F. Tekola-Ayele, L. Pagani, I. Tach-mazidou, et al., 2015 The african genome variation project shapes medical genetics in africa. Nature 517: 327–332.

Gutenkunst, R. N., R. D. Hernandez, S. H. Williamson, and C. D. Bustamante, 2009 Inferring the joint demographic history of multiple populations from multidimensional SNP frequency data. PLoS Genet 5: e1000695.

Haak, W., I. Lazaridis, N. Patterson, N. Rohland, S. Mallick, et al., 2015 Massive migration from the steppe was a source for indo-european languages in europe. Nature 522: 207.

Henn, B. M., T. E. Steele, and T. D. Weaver, 2018 Clarifying distinct models of modern human origins in africa. Current Opinion in Genetics & Development 53: 148–156, Genetics of Human Origins.

Hollfelder, N., C. M. Schlebusch, T. Günther, H. Babiker, H. Y. Hassan, et al., 2017 Northeast african genomic variation shaped by the continuity of indigenous groups and eurasian migrations. PLOS Genetics 13: 1–17.

Hudson, R., 2002 Generating samples under a Wright–Fisher neutral model of genetic variation. Bioinformatics 18: 337–338, doi:10.1093/bioinformatics/18.2.337.

Kelleher, J., Y. Wong, A. W. Wohns, C. Fadil, P. K. Al-bers, et al., 2019 Inferring whole-genome histories in large population datasets. Nature Genetics 51: 1330–1338, doi.org/10.1038/s41588-019-0483-y.

Kuhlwilm, M., I. Gronau, M. J. Hubisz, C. de Filippo, J. Prado-Martinez, et al., 2016 Ancient gene flow from early modern humans into eastern neanderthals. Nature 530: 429–433.

Li, H. and R. Durbin, 2011 Inference of human population history from individual whole-genome sequences. Nature 475: 493–496, doi:10.1038/nature10231.

Llorente, M. G., E. R. Jones, A. Eriksson, V. Siska, K. W. Arthur, et al., 2015 Ancient ethiopian genome reveals extensive eurasian admixture in eastern africa. Science 350: 820–822.

Lohse, K., M. Chmelik, S. H. Martin, and N. H. Barton, 2016 Efficient Strategies for Calculating Blockwise Likelihoods Under the Coalescent. Genetics 202: 775+.

Lohse, K., R. J. Harrison, and N. H. Barton, 2011 A General Method for Calculating Likelihoods Under the Coalescent Process. Genetics 189: 977–U398.

Mazet, O., W. Rodriguez, S. Grusea, S. Boitard, and L. Chikhi, 2016 On the importance of being structured: instantaneous coalescence rates and human evolution–lessons for ances-tral population size inference? Heredity 116: 362–371, doi:10.1038/hdy.2015.104.

Meyer, M., M. Kircher, M.-T. Gansauge, H. Li, F. Racimo, et al., 2012 A high-coverage genome sequence from an archaic denisovan individual. Science 338: 222–226.

Moorjani, P., Z. Gao, and M. Przeworski, 2016 Human Germline Mutation and the Erratic Evolutionary Clock. PLoS Biol 14: e2000744, doi:10.1371/journal.pbio.2000744.

Orozco, P., 2016 The devil is in the details: the effect of pop-ulation structure on demographic inference. Heredity 116: 349–350, doi:10.1038/hdy.2016.9.

Patin, E., M. Lopez, R. Grollemund, P. Verdu, C. Harmant, et al., 2017 Dispersals and genetic adaptation of bantu-speaking populations in africa and north america. Science 356: 543–546.

Pickrell, J. K., N. Patterson, C. Barbieri, F. Berthold, L. Gerlach, et al., 2012 The genetic prehistory of southern Africa. Nature Communications 3: 1143

Pickrell, J. K., N. Patterson, P.-R. Loh, M. Lipson, B. Berger, et al., 2014 Ancient west eurasian ancestry in southern and eastern africa. Proceedings of the National Academy of Sciences 111: 2632–2637.

Prüfer, K., F. Racimo, N. Patterson, F. Jay, S. Sankararaman, et al., 2014 The complete genome sequence of a neanderthal from the altai mountains. Nature 505: 43–49.

Pudlo, P., J. Marin, A. Estoup, J. Cornuet, M. Gautier, et al., 2016 Reliable abc model choice via random forests. Bioinformatics 32: 859–866.

Raghavan, M., M. Steinrücken, K. Harris, S. Schiffels, S. Ras-mussen, et al., 2015 Genomic evidence for the pleistocene and recent population history of native americans. Science 349.

Rogers, A. and R. Bohlender, 2014 Bias in estimators of archaic admixture. Theoretical Population Biology 100: 63–78, doi.org/10.1016/j.tpb.2014.12.006.

Rogers, A. R., R. J. Bohlender, and C. D. Huff, 2017 Early history of neanderthals and denisovans. Proceedings of the National Academy of Sciences 114: 9859–9863.

Scally, A. and R. Durbin, 2012 Revising the human mutation rate: implications for understanding human evolution. Nature Reviews Genetics 13: 745–753.

Scerri, E. M., M. G. Thomas, A. Manica, P. Gunz, J. T. Stock, et al., 2018 Did our species evolve in subdivided populations across africa, and why does it matter? Trends in Ecology & Evolution 33: 582–594.

Schiffels, S. and R. Durbin, 2014 Inferring human population size and separation history from multiple genome sequences. Nature Genetics 46: 919–925, doi:10.1038/ng.3015.

Schlebusch, C. M. and M. Jakobsson, 2018 Tales of human mi-gration, admixture, and selection in africa. Annual Review of Genomics and Human Genetics 19: null, PMID: 29727585.

Schlebusch, C. M., H. Malmström, T. Günther, P. Sjödin, A. Coutinho, et al., 2017 Southern african ancient genomes es-timate modern human divergence to 350,000 to 260,000 years ago. Science 358: 652–655.

Schlebusch, C. M., P. Skoglund, P. Sjödin, L. M. Gattepaille, D. Hernandez, et al., 2012 Genomic variation in seven khoe-san groups reveals adaptation and complex african history. Science 338: 374–379.

Schraiber, J. G. and J. M. Akey, 2015 Methods and models for unravelling human evolutionary history. Nature Reviews Genetics 16: 727–740.

Skoglund, P., A. Götherström, and M. Jakobsson, 2011 Estimation of population divergence times from non-overlapping genomic sequences: Examples from dogs and wolves. Molecular Biology and Evolution 28: 1505–1517.

Skoglund, P., S. Mallick, M. C. Bortolini, N. Chennagiri, T. Hüne-meier, et al., 2015 Genetic evidence for two founding popula-tions of the americas. Nature 525.

Skoglund, P., P. Sjödin, T. Skoglund, M. Lascoux, and M. Jakobsson, 2014 Investigating population history using temporal genetic differentiation. Mol Biol Evol 31: 2516–2527, doi:10.1093/molbev/msu192.

Slatkin, M., 1996 Gene genealogies within mutant allelic classes. Genetics 143: 579–587.

Speidel, L., M. Forest, S. Shi, and S. Myers, 2019 A method for genome-wide genealogy estimation for thousands of samples. Nature Genetics 51: 1321–1329.

Stringer, C., 2016 The origin and evolution of *Homo sapiens*. Philosophical Transactions of the Royal Society B: Biological Sciences 371: 20150237.

Tavaré, S., D. J. Balding, R. C. Griffiths, and P. Donnelly, 1997 Inferring coalescence times from DNA sequence data. Genetics 145: 505–518.

Terhorst, J., J. A. Kamm, and Y. S. Song, 2017 Robust and scalable inference of population history from hundreds of unphased whole-genomes. Nature genetics 49: 303–309.

Theunert, C. and M. Slatkin, 2018 Estimation of population divergence times from snp data and a test for treeness. bioRxiv.

Triska, P., P. Soares, E. Patin, V. Fernandes, V. Cerny, et al., 2015 Extensive Admixture and Selective Pressure Across the Sahel Belt. Genome Biology and Evolution 7: 3484–3495.

Černý, V., I. Kulichová, E. S. Poloni, J. M. Nunes, L. Pereira, et al., 2018 Genetic history of the african sahelian populations. HLA 91: 153–166.

Wakeley, J., 2009 Coalescent Theory: An Introduction. Roberts & Company Publishers, Greenswood Village, Colorado, first edition.

Wakeley, J. and J. Hey, 1997 Estimating ancestral population parameters. Genetics 145: 847–855.

Wald, A., 1949 Note on the consistency of the maximum like-lihood estimate. The Annals of Mathematical Statistics 20: 595–601.

Wang, K., I. Mathieson, J. O’Connell, and S. Schiffels, 2020 Tracking human population structure through time from whole genome sequences. PLOS Genetics 16: 1–24.

Wilkinson-Herbots, H. M., 2008 The distribution of the coales-cence time and the number of pairwise nucleotide differences in the “isolation with migration” model. Theoretical Population Biology 73: 277–288.

